# The LOTUS domain of Oskar promotes localisation of both protein and mRNA components of *Drosophila* germ plasm

**DOI:** 10.1101/2025.05.02.651258

**Authors:** Anastasia Repouliou, John R. Srouji, Emily L. Rivard, Andrés E. Leschziner, Cassandra G. Extavour

## Abstract

Germ cells transmit genetic information to the next generation in multicellular organisms. In *Drosophila melanogaster*, germ cells are determined by germ plasm, a specialised cytoplasm assembled by the Oskar protein. The current view of the molecular mechanism of germ plasm assembly attributes recruitment of protein and mRNA germ plasm components to distinct domains of the Oskar protein, called the LOTUS and OSK domains respectively. However, most evidence for this model is based on *in vitro* studies. Here we test the ability of Oskar variants to assemble functional germ plasm *in vivo*. We found that Vasa recruitment was largely unperturbed by LOTUS deletion or mutations *in vivo*. In contrast, *nanos* and *pgc* mRNA recruitment was affected by LOTUS domain perturbations, despite the current model attributing mRNA recruitment to the distinct OSK domain. Taken together, these data suggest a revision of the prevailing modular view of Oskar’s structure-function mechanism.

## INTRODUCTION

Patterning of a developing animal embryo is tightly spatiotemporally regulated. Premature or ectopic activation of the molecular factors specifying cell fate can lead to severe developmental defects. Germ line fate is particularly important because the eggs and sperm are uniquely capable of transmitting genetic information across generations. In *Drosophila melanogaster*, germ line specification requires the coalescence of multiple fate-determining elements at the posterior of the oocyte ^1–5^. The assembly of this set of molecules, collectively referred to as “germ plasm” ^6,7^ is orchestrated by the Oskar protein ^8^. Oskar is necessary for germ plasm accumulation and localisation ^9^, and subsequently critical for formation of unique cells at the posterior of the embryo called pole cells, which are the primordial germ cells of this animal ^8–13^. Despite the importance of Oskar in germ line specification, a comprehensive *in vivo* molecular mechanism for Oskar-mediated germ plasm assembly is lacking.

During early oogenesis, *oskar* mRNA is maternally produced ^11^ and loaded into the oocyte as part of a translationally repressed ribonucleoprotein complex ^8,11,12,14,15^. Once accumulated in the oocyte, *oskar* is tightly localised to the posterior ^11,12^, where the transcript is translated ^16^, producing two isoforms from alternate translation start sites ^9^. The protein products of *oskar* go on to assemble and localise germ plasm components. Germ plasm then becomes engulfed in the first cells that form in the developing *D. melanogaster* embryo from the posterior-most nuclei of the embryonic syncytium ^11^. These “pole cells” will adopt germ cell fate and constitute the germ line of the future fly. Oskar is required for both germ plasm assembly and pole cell formation ^9^.

The Oskar protein sequence is traditionally divided into four domains for functional and evolutionary study (Fig. 1A): an N-terminal disordered extension, present only in the long isoform of Oskar (residues 1-138), the LOTUS domain (residues 139-240), a disordered linker (residues 241-400), and the OSK domain (residues 401-606). The folded LOTUS and OSK domains play key roles in Oskar function, but precisely how they contribute to the intermolecular interactions underlying germ plasm assembly *in vivo* have not been definitively determined. Investigating the degree to which the distinct domains of Oskar function independently or interdependently is a critical step towards understanding the molecular mechanism of Oskar-mediated germ plasm assembly.

**Figure 1.**
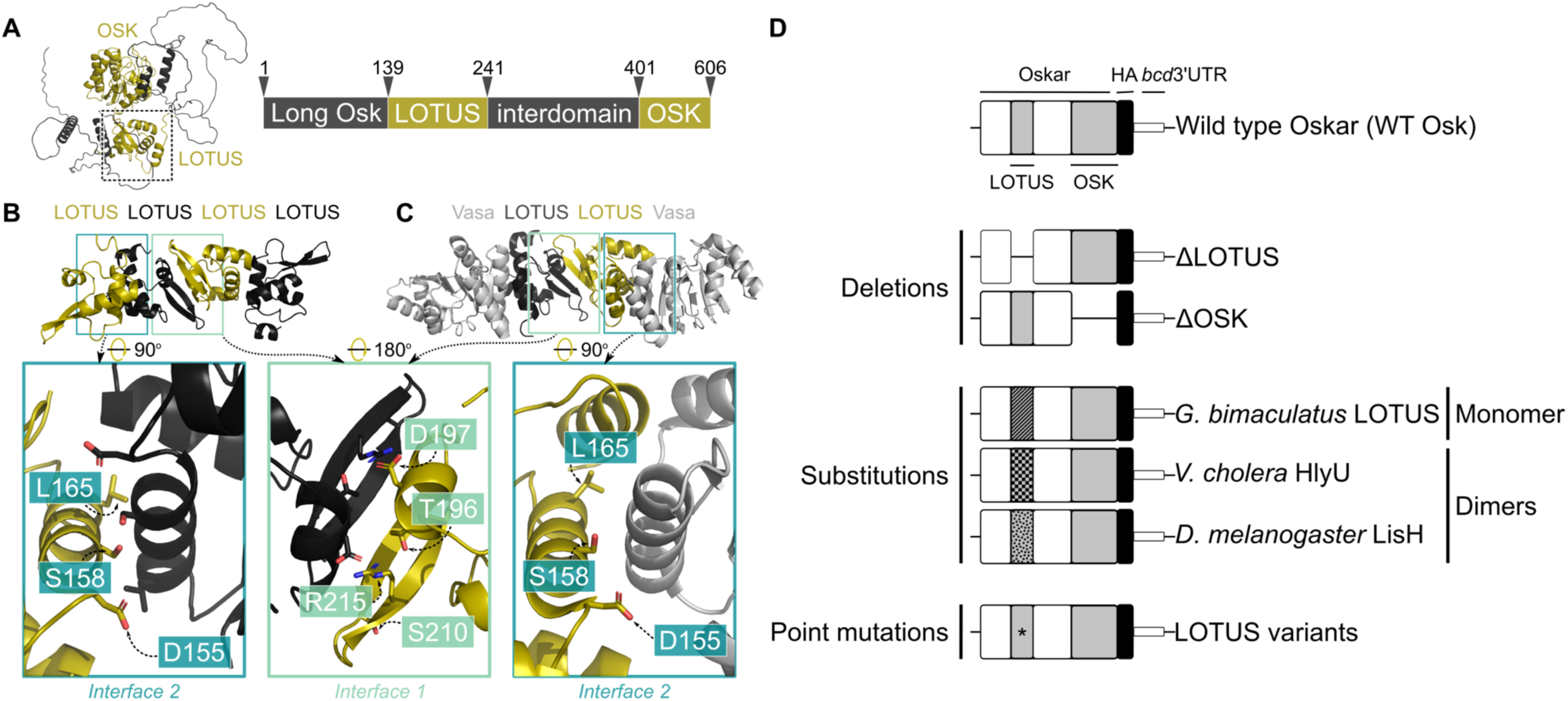
Using predicted molecular interactions of the Oskar LOTUS domain to inform *in vivo* experimental design. **(A)** Left: structure of full-length *D. melanogaster* Oskar as predicted by AlphaFold2 ^68,69^. Folded OSK and LOTUS domains highlighted in gold. LOTUS domain outlined in dashed box. Right: schematic representation of Oskar domains. Arrows and numbered amino acids indicate boundaries between domains. **(B)** Crystal structure of LOTUS tetramer. **(C)** Crystal structure of LOTUS dimer in complex with two Vasa monomers (RCSB PDB 5NT7, Jeske et al. 2017 ^36^). In (B) and (C), interface 1, mediating predicted LOTUS dimerisation, is outlined in green. Interface 2, corresponding to an alternative putative LOTUS dimerisation interface in (B) and to the LOTUS-Vasa binding interface in (C), is outlined in blue. In the magnified insets below (B) and (C), the residues involved in predicted LOTUS dimerisation and LOTUS-Vasa binding are shown as sticks colored in red and labelled in the gold protomer. **(D)** To investigate the role of LOTUS dimerisation and Vasa-binding *in vivo*, we designed UAS-controlled *D. melanogaster oskar* variant cDNA constructs. All constructs contain a C-terminal HA-tag and the *bicoid* 3’UTR (black) to target the transcript to the oocyte anterior and allow detection of the translated variant protein. Oskar variants investigated were wild type Oskar (WT Osk, positive control), deletions of the entire LOTUS and OSK domains, substitutions of LOTUS domain with *G. bimaculatus* Oskar LOTUS domain (striped), found to be monomeric (Fig. S1), with dimeric *Vibrio cholera* winged helix-turn-helix HlyU (checkered), and with dimeric *D. melanogaster* Lis1 LisH domain (polka- dotted), and point mutations in the residues labelled in (B) and (C).

The N-terminal LOTUS (<u>L</u>imkain, <u>O</u>skar and <u>TU</u>dor containing protein <u>S</u> 5 and 7) or OST- HTH (<u>OS</u>kar-<u>T</u>DRD5/TDRD7 <u>H</u>elix-<u>T</u>urn-<u>H</u>elix) domain ^17,18^ (Pfam ^19^ family PF12872) folds into a winged helix-turn-helix (wHTH) ^20^conformation ^21,22^. Members of this structurally related protein family commonly participate in nucleic acid binding ^23–25^, including the HlyU transcription factor from *Vibrio cholera* ^26,27^, the SarR transcription factor from *Staphylococcus aureus* ^28^, and components of ribonucleoprotein complexes ^29,30^. LOTUS family members can also participate in protein-protein interactions, such as the C-terminal homodimerisation module of *Saccharomyces cerevisiae* Sir3 ^31^. The LOTUS domain is also found in some other germ plasm protein components ^17,20^, including the RNA-binding Tudor 5 and Tudor 7 proteins ^30^. Because germ plasm contains multiple specific mRNAs, LOTUS was originally hypothesised to promote RNA binding ^17,20^, but no direct evidence has supported this hypothesis to date. Instead, biochemical studies support the hypothesis that the C-terminal OSK domain ^32–35^ directly binds RNA ^22,36^: OSK binds the 3’-UTR of *nanos* and *oskar* mRNAs *in vitro* ^22^ and full-length *nanos*, *polar granule component* (*pgc*), and *germ cell-less* (*gcl*) mRNAs when UV-crosslinked and immunoprecipitated from early-stage embryos ^36^. The prevailing model thus attributes mRNA recruitment to the OSK domain.

Germ plasm also contains specific proteins, of which Vasa comprises a heavily examined example ^10,13,37^. Strong evidence supports a direct Oskar-Vasa binding interaction ^21,36,38–41^. However, investigations into exactly how Oskar interacts with Vasa have yielded contradictory findings: a yeast two-hybrid assay showed Oskar-Vasa binding was strongest for the C-terminal end of the protein, which contains the OSK domain ^39^. However, while necessary, OSK was not by itself sufficient for Vasa binding *in vitro* ^39^. Later biochemical and structural work showed that the LOTUS domain binds and activates Vasa *in vitro* ^21,36^. Given this latest evidence ^21,36^, the prevailing model posits that Vasa recruitment is mediated by the LOTUS domain.

While *in vitro* characterisations continue to generate useful hypotheses regarding Oskar function, a rigorous probe of the interactions between Oskar and germ plasm components *in vivo* is required to develop a biologically relevant molecular mechanistic model for Oskar function. Here, we elucidate the role of the LOTUS domain in recruiting functional germ plasm. We use structural and biochemical assays to recapitulate and validate observed LOTUS-Vasa and LOTUS homodimerisation interactions *in vitro*, and developmental genetics approaches to test the effect of LOTUS genetic perturbations on the accumulation of germ plasm components and the assembly of functional germ plasm *in vivo*. Our findings show that pole cell formation is susceptible to perturbations in the LOTUS domain. However, the lower incidence of pole cell formation in LOTUS variants does not seem attributable to impaired Vasa recruitment, which appears robust to disruptions of the Vasa-LOTUS binding interface. Instead, LOTUS perturbations result in deficient *nanos* and *pgc* mRNA recruitment. Our findings therefore implicate LOTUS in germ plasm mRNA recruitment, with dimerisation appearing to be at least one aspect of its recruitment mechanism. Overall, our results suggest a new, holistic model for germ plasm nucleator function, contrasting with the prevailing modular view of Oskar’s function and shedding new light on the molecular mechanism of Oskar-mediated germ plasm assembly.

## RESULTS

### The Oskar LOTUS domain forms homodimers in vitro

To investigate the molecular mechanism of Oskar *in vivo*, we used published and novel biochemical and structural characterisations of the Oskar LOTUS domain to generate testable predictions about the specific molecular interactions that could be mediating Oskar-mediated germ plasm assembly *in vivo*. Published *in vitro* studies ^21,22^ suggested that purified Oskar LOTUS domain behaves as a dimer based on size exclusion chromatography (SEC) ^21^, static light scattering (SLS) ^21^, and multi-angle light scattering (MALS) ^22^. Consistent with these reports, we found that the LOTUS domain forms dimers *in vitro* using analytical SEC ^42^ (Fig. S1), MALS ^43^, and bacterial two-hybrid analysis ^44,45^. In analytical SEC across various ionic strengths, we observed a retention volume of the fly *D. melanogaster* LOTUS domain that was lower than expected for a monomer (12.3 kDa) and the apparent molecular weight (MW) was close to that of a dimer (Fig. S1). In contrast, the cricket Oskar LOTUS domain of the *Gryllus bimaculatus* eluted approximately as expected for a monomer (10.5 kDa) across all ionic strengths (Fig. S1A). SEC-MALS ^43,46^ suggested that the fly LOTUS domain was likely to form a dimer (MW = 21.55 ± 4.13 kDa) and that the cricket LOTUS domain was monomeric (MW = 10.03 ± 3.89 kDa) in solution (data not shown). Finally, we confirmed that the fly LOTUS domain oligomerises and the cricket LOTUS domain does not, in a bacterial two-hybrid assay ^44,45^ (BACTH system, Euromedex; data not shown).

Published crystal structures of the LOTUS domain ^21,22^ reveal the details of the dimerisation implied by the biochemical assays detected above. We independently obtained crystals of the LOTUS domain (residues 139-241) that diffracted to ∼3.2Å (P4 - form 1), ∼3.1Å (I4 - form 2), and ∼4.2Å (C222 - form 3; Table S1). Because none of these diffraction datasets were soluble by molecular replacement, we performed random sitting drop vapor diffusion screening with HgCl2-derivatised LOTUS, obtaining a crystal with diffraction up to 2.0Å (P1211 - form 4; Table S1) which we were able to solve with molecular replacement. We used the solved form 4 structure to successfully phase the native diffraction data collected from crystal forms 1-3 via molecular replacement. Our structures (Fig. 1B) corroborated the main features of the published LOTUS structures (Fig. 1C).

Despite solving crystals originating from different space groups, both previously published studies ^21,22^ as well as our own work, reveal two potential homodimerisation interfaces, mainly stabilised through hydrogen bonds. Interface 1 (Fig. 1B-C), common in all forms, consists of a completed β-sheet, mediated by the alignment of the β-hairpins from both protomers ^21,22^. The dimer is stabilised by salt bridges across the interface between the side chains of D197 from one protomer, and T196 and R216 from the other, burying hydrophobic side chains of L200, A207, and F217 ^21,22^. Interface 2 (Fig. 1B-C), also seen in some of the previously reported crystal forms ^21,22^, is formed via helices α2 and α4 of participating protomers, mediated by hydrophilic interactions and the seclusion of hydrophobic side chains, involving residues D155, S158, R161, A162, L165, and L184. Interface 2, but not interface 1, is commonly observed among dimers of unrelated wHTH domain dimers ^23,25^. Interestingly, interface 2 partially overlaps with the LOTUS domain interface shown to directly contact Vasa in a previously reported co-crystal ^36^ (Fig. 1C).

To determine the impact *in vitro* of mutations in residues at the predicted dimerisation interfaces evident in the crystal structures (Fig 1B-C), we applied BACTH and analytical SEC to LOTUS domain variants containing point mutations at interfaces 1 and 2 and determined their oligomeric state. We found that interface 2 mutations did not perturb oligomerisation *in vitro* (Fig. S1D). In contrast, most interface 1 mutations resulted in insoluble protein (precluding analytical SEC experiments) and only T196S and the triple mutant showed detectable *in vitro* oligomerisation using BACTH (Fig. S1D). Of the interface 1 mutants that yielded soluble protein (T196S, R215E, and R215Q), R215E showed disrupted dimerisation *in vitro* (Fig. S1D). The monomeric state of R215E is consistent with published analytical gel filtration and SLS ^21^ and presumed to be the result of interference with the R215-D197 interaction ^21,22^. R215Q exhibited a temperature-dependent effect on oligomerisation, showing detectable oligomerisation via BACTH at room temperature but not at 30°C (Fig. S1D). Furthermore, S210P elicited no oligomerisation signal via BACTH, consistent with published SEC-MALS ^22^, which was interpreted as the result of a kink disrupting the β-strand conformation ^22^.

*LOTUS deletions, substitutions, and point mutations decrease the incidence of pole cell formation* To evaluate the functional biological relevance of the identified dimerisation and Vasa-binding interfaces *in vivo*, we exploited the fact that ectopically localised *oskar* is sufficient for the accumulation of active germ plasm and subsequent pole cell specification ^10,13^. Published experiments demonstrated this sufficiency by targeting the *oskar* cDNA to the oocyte anterior via fusion to the 3’-UTR of the anterior morphogen *bicoid* (*bcd*) ^10^. The anteriorly-localised *oskar* transcript is translated and the resulting ectopic Oskar protein can localise known germ plasm components, including the Vasa and Nanos proteins ^10,13,47^, and produce ectopic pole cells at the anterior ^13^. Capitalising on this finding, we used the ΦC31 system ^48^ to generate transgenic *oskar* constructs encoding HA-tagged *oskar* targeted to the oocyte anterior via fusion to the *bcd* 3’-UTR (Fig. 1D), under UAS control. We used a maternal α-tubulin GAL4 driver ^49^ to express the *oskar* constructs in the female germ line during oogenesis, when Oskar is actively assembling germ plasm. Each construct carried a specific genetic perturbation aimed at elucidating the functional relevance of the LOTUS domain and its dimerisation and Vasa-binding interfaces *in vivo* (Fig. 1D).

To investigate the function of the folded domains of Oskar we generated LOTUS and OSK domain deletions (abbreviated as ΔLOTUS and ΔOSK, Fig. 1D). To evaluate the relevance of LOTUS dimerisation, we substituted the LOTUS domain with the related but monomeric cricket LOTUS domain ^50^ (abbreviated as Gb LOTUS) or with either of two exogenous dimerisation domains: the *Vibrio cholera* wHTH motif HlyU (UniProt Acc. No. C3LST3) residues 1-109 ^27^ (abbreviated as Vc HlyU), and the *D. melanogaster* LisH domain of the dynein regulator lis1 (UniProt Acc. No. Q7KNS3) residues 1-86 ^27,51^, (abbreviated as Dm LisH, Fig. 1D). There is precedent for HlyU restoring dimerisation and protein function when put in place of an endogenous wHTH motif ^31^. Finally, to elucidate the functional relevance of the homodimerisation and Vasa- binding interfaces seen in the crystal structures, we introduced point mutations at relevant residues in the two interfaces (Fig. 1D). For simplicity, all oocytes with the Oskar variant-HA-*bcd*3’UTR genotype, as well as embryos from mothers with the Oskar variant-HA-*bcd*3’UTR genotype, will be referred to herein by the variant itself, be it the deleted domain (e.g. ΔLOTUS), the substituted domain (e.g. Gb LOTUS), or the point mutation (e.g. D155A).

We assessed the ability of the anteriorly-targeted Oskar variants to assemble functional germ plasm by measuring the incidence of anterior pole cell formation and the number of pole cells formed in stage 5 embryos from mothers carrying the genetic perturbations (Fig. 2A-B). Embryos that displayed weakly-budding cells, Vasa-negative budding cells, or non-budding cells with spherical nuclei were scored as “anterior pole cell attempts” (Fig. 2A, Fig. S2). Embryos of our wild type strain (Oregon R) have an average of 39.6 pole cells at the posterior (Fig. 2C), and targeting wild type *oskar* to the anterior generates an average of 20.5 ectopic anterior pole cells (Fig. 2C) in 15.1% of embryos (Table S2). Eight of 14 tested *oskar* variants were unable to generate any ectopic anterior pole cells at all (Fig. 2B; Table S2). The six variants that did, generated the same average number of ectopic anterior pole cells as controls (p > 0.05 for all cases except D197N (p = 0.005) and S210P (p=5.4e-15, but note that these cases had only 1/307 and 2/355 embryos with anterior pole cells, and that statistical tests have very limited power with such small sample sizes; Fig. 2C, Table S3), but with significantly lower penetrance, ranging from 0.1% (ΔLOTUS; p = 3.00e-11) to a maximum of 2.1% (T196S; p = 1.09e-09; Fig. 2B, Table S2-3). These results suggest that the LOTUS domain is important for germ plasm function, but also that even in its complete absence (ΔLOTUS), functional germ plasm can still form, albeit at very low incidence.

**Figure 2.**
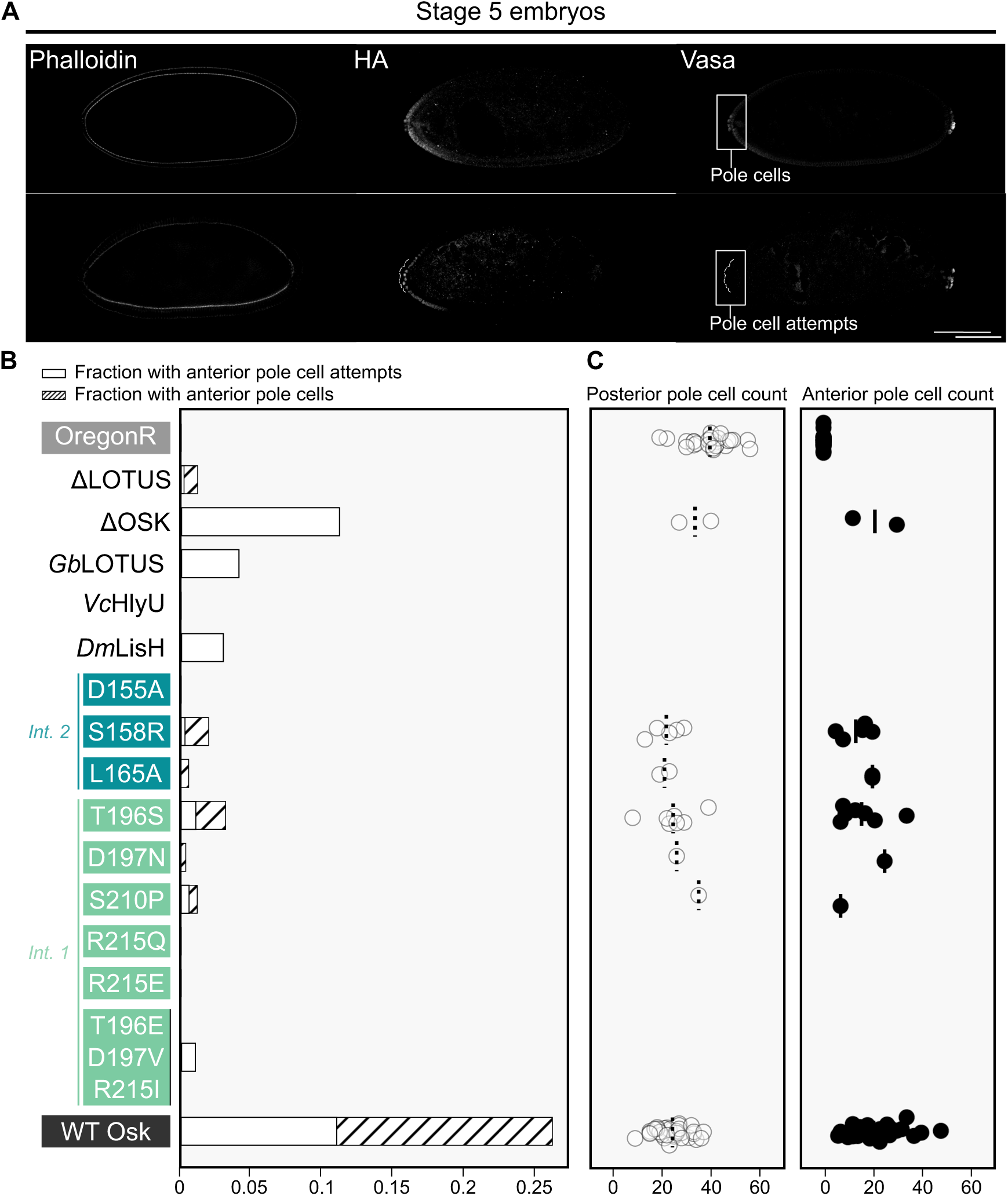
Ectopic pole cell induction is decreased by all tested perturbations in the LOTUS domain. **(A)** Optical sections of confocal micrographs of stage 5 embryos from *y*^1^ *v^1^/w*; P{y^+^ v^+^ = (oskar variant-HA-bcd3’UTR)}attp40/+; P{w^+mC^=matalpha4-GAL-VP16}V37/+* mothers. Fixed embryos were stained for nuclei (DAPI, not shown), F-actin (phalloidin) for staging, the HA epitope, and Vasa (anti-Vasa antibody) to mark pole cells. Anterior is to the left. Micrographs selected to show an example of successful anterior pole cells (top) and an example of pole cell attempts (bottom, dashed box, anterior buds outlined in white, background subtracted for visualisation purposes). See Fig. S2 for more examples of pole cell attempts. Scale bar = 100μm and applies to all panels. **(B)** Stacked bar plot of fraction of collected embryos with anterior pole cell attempts (white) and with successfully formed anterior pole cells (striped). **(C)** Counts of posterior (left) and anterior (right) pole cells for variants with a nonzero fraction of embryos with anterior pole cells and a random subset of Oregon R embryos. Mean pole cell count indicated by dashed (left) or solid (right) black line.

### Vasa recruitment in vivo is largely robust to LOTUS genetic perturbations

We hypothesised that deficient pole cell formation could result from insufficient or defective recruitment of germ plasm components. Given the strong *in vitro* links between the Oskar LOTUS domain and Vasa protein, we asked whether the genetic perturbations to the LOTUS domain affected Vasa enrichment at the anterior of stage 10 oocytes and stage 1-2 embryos (Fig. 3A). We quantified Vasa enrichment as the integral of the z-scores in the anterior-most 15% of the oocyte or embryo (Fig. 3B, D). We also quantified HA-Vasa colocalisation at the oocyte and embryo anterior as the Pearson correlation between the HA and Vasa signal intensity in a given anterior pixel with the 17.5% highest HA intensity values for each micrograph (Fig. S3; see Methods).

**Figure 3.**
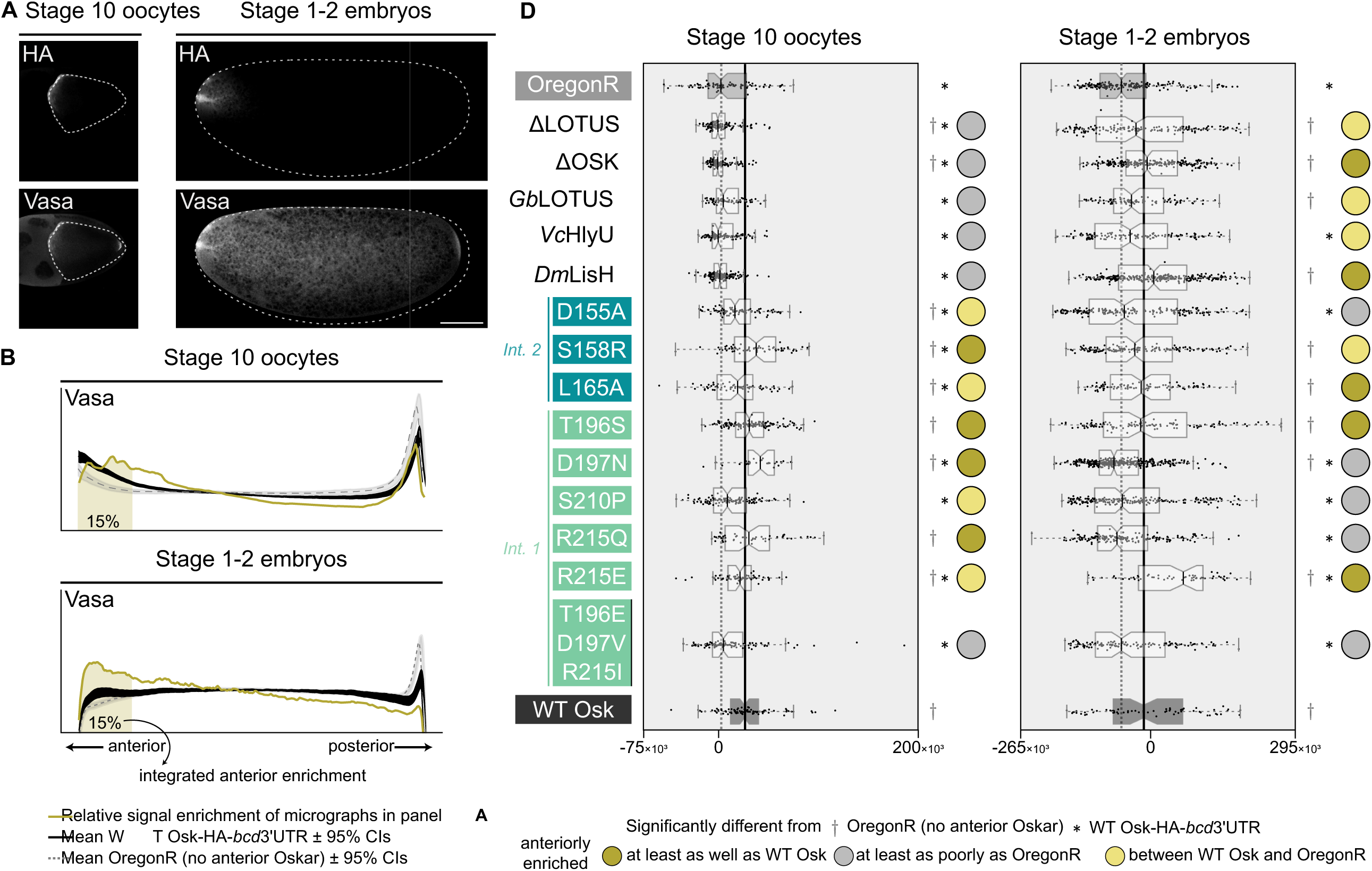
Vasa recruitment *in vivo* is robust to perturbations in the LOTUS domain. **(A)** Optical sections of confocal micrographs of stage 1-2 embryos of stage 10 oocytes and stage 1-2 embryos from *y^1^ v^1^/w*; P{y^+^ v^+^ = (oskar variant-HA-bcd3’UTR)}attp40/+; P{w^+mC^=matalpha4- GAL-VP16}V37/+* mothers, stained as described in Figure 2 and selected to show an example of successful anterior enrichment of Vasa. Anterior to the left. Scale bar = 50μm and applies to all panels. **(B)** Relative Vasa signal enrichment (z-score) for the micrographs in (A) (gold), mean z- score lines and 95% confidence intervals for *y^1^ v^1^/w*; P{y^+^ v^+^ = (wt-osk-HA-bcd3’UTR)}attp40/+; P{w^+mC^=matalpha4-GAL-VP16}V37/+* (black line, positive control) and Oregon R (grey line, negative control). **(C)** Box plots of integrated anterior enrichment (integral of 15% anterior-most z-score values) for Vasa in stage 10 oocytes (left) and stage 1-2 embryos (right). Oregon R (negative control) in grey, median as grey dashed line. *y^1^ v^1^/w*; P{y^+^ v^+^ = (wt-osk-HA- bcd3’UTR)}attp40/+; P{w^+mC^=matalpha4-GAL-VP16}V37/+* (positive control) in black, median as solid black line. Significant difference from positive and negative control distributions (p-value (< 0.05) estimated using a bootstrap resampling test (10,000 iterations), comparing the observed difference in means between sample and control to a null distribution generated by resampling with replacement) indicated by asterisk and cross, respectively. Circles indicate interpretation of data: mRNA integrated anterior enrichment in variant significantly higher than negative control (dark golden), between positive and negative control (light golden), or significantly lower than positive control (grey).

In both oocytes and early embryos, Vasa anterior enrichment was largely unperturbed (Fig. 3C). The LOTUS and OSK domain deletions and the LOTUS domain substitutions failed to anteriorly enrich Vasa in late-stage oocytes (Fig. 3C). However, all except the Vc HlyU variant recovered anterior Vasa enrichment to the same degree as the control by the early embryonic stage (Fig. 3C). In contrast, the LOTUS domain point mutations were able to drive control or near-control levels of anterior Vasa enrichment in stage 10 oocytes, the triple mutant (T196E-D197V- R215I) being the only exception (Fig. 3C). However, half of the point mutants (D155A from interface 2, D197N, S210P, and R215Q from interface 1) were unable to maintain this Vasa anterior enrichment through to the embryonic stage (Fig. 3C). Neither the interface 2 mutants (D155A, S158R, and L165A) nor interface 1 mutants D197N and R215E completely abolished anterior Vasa enrichment *in vivo* (Fig. 3C, S3), even though these mutants disrupted LOTUS-Vasa binding *in vitro* (Fig. S1D). Three of five point mutations at predicted dimerisation interface 1 (D197N, S210P and R215Q), as well as the triple mutant at this interface (T196E-D197V-R215I) failed to maintain control levels of anterior Vasa, while one of three mutations at interface 2 (D155A) failed to do so (Fig. 3C). This suggests that of these two predicted Oskar dimer conformations, dimerisation at interface 1 may be more relevant to localising Vasa to germ plasm *in vivo*.

If Vasa is recruited to germ plasm through direct interaction with Oskar via the LOTUS domain, then we would predict that the degree of colocalisation of Oskar (quantified in our design by detection of the HA tag placed on each variant) and Vasa proteins would correlate with the enrichment of Vasa described above for each mutant. We tested this prediction by determining the Pearson correlation between the HA and Vasa signal intensities of the anterior pixels with the 17.5% highest HA intensity values (Fig. S3). We found that all perturbations showing lower Vasa anterior enrichment than the positive control showed HA-Vasa colocalisation comparable to or stronger than that seen in WT Osk (at the oocyte stage: βLOTUS, βOSK, LOTUS substitutions, triple mutant; at the embryonic stage: D155A, D197N, S210P, R215Q, triple mutant; Fig. 3C, Fig. S3). This could mean that loss of Vasa enrichment in these LOTUS domain variants *in vivo* may not be due to significantly impaired interactions between Oskar and Vasa proteins, but rather to an indirect interaction between Oskar and other germ plasm components. In all, Vasa recruitment by Oskar is not severely disrupted by LOTUS domain deletion, substitutions, or point mutations.

### The LOTUS domain is critical for the recruitment in vivo of nanos and pgc mRNAs

While Vasa protein recruitment appears largely robust to perturbations in the LOTUS domain, all tested LOTUS variants either failed to induce the formation of ectopic pole cells, or did so very inefficiently (Fig. 2B, Table S2). Although a direct role in germ plasm mRNA binding has not been reported for the Oskar LOTUS domain, LOTUS domains do perform RNA binding functions in other proteins ^17,52^, and Oskar’s LOTUS domain can bind G4 RNA^53^ in an ELISA assay^52^. We therefore asked whether disruption of the LOTUS domain affected *in vivo* enrichment of two critical germ plasm mRNA components, *nanos* and *polar granule component* (*pgc*), at the anterior of stage 1-2 embryos (Fig. 4A). We measured *nanos* and *pgc* integrated anterior enrichment (Fig. 4B-C) as well as HA-*nanos* and HA-*pgc* colocalisation as the Pearson correlation (Fig. S4), as described above.

**Figure 4.**
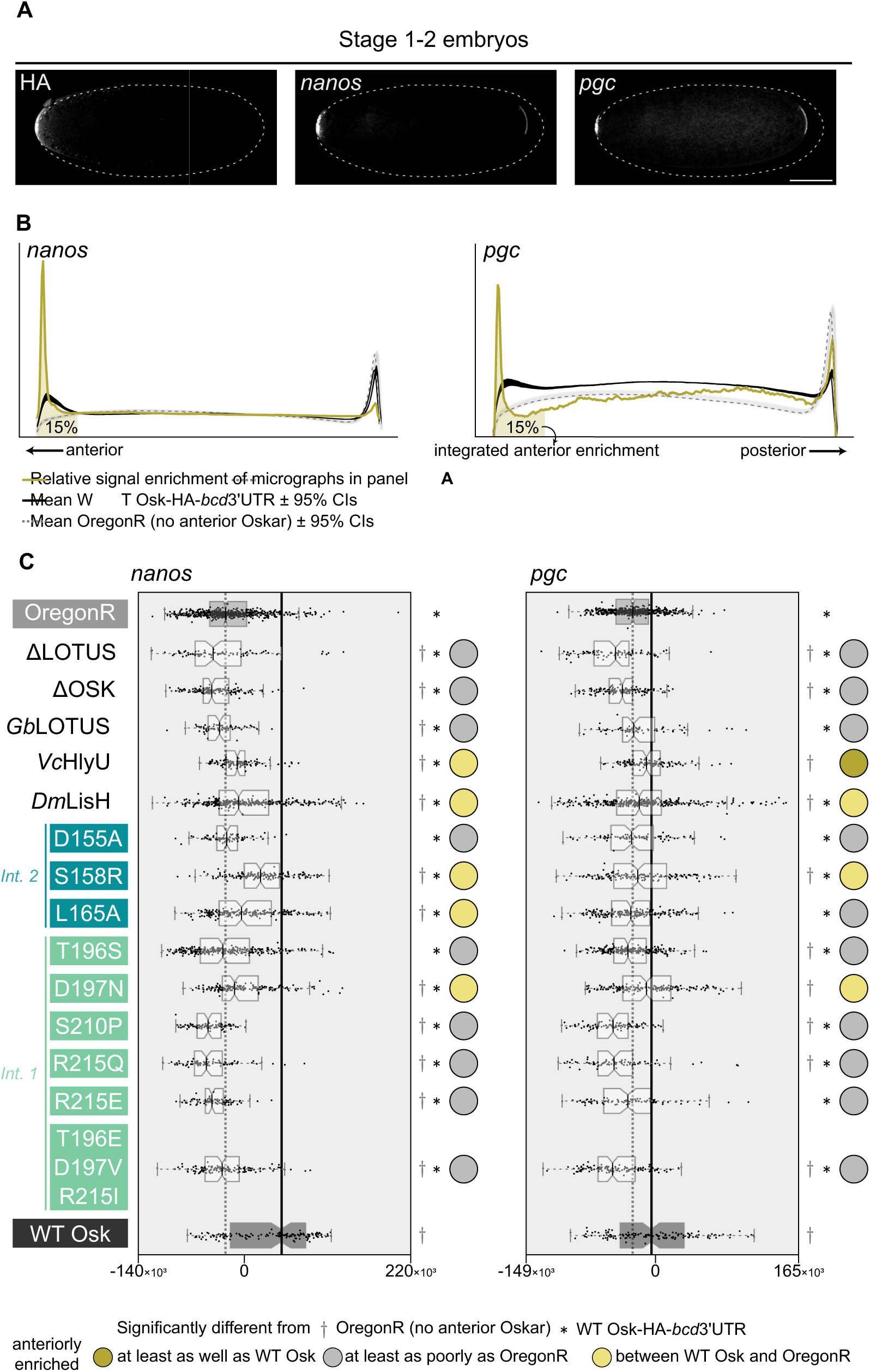
The LOTUS domain is involved in *nanos* and *pgc* recruitment. **(A)** Optical sections of confocal micrographs of stage 1-2 embryos from *y^1^ v^1^/w*; P{y^+^ v^+^ = (oskar variant-HA- bcd3’UTR)}attp40/+; P{w^+mC^=matalpha4-GAL-VP16}V37/+* mothers stained as described in Figure 2 and selected to show an example of successful anterior enrichment of *nanos* and *pgc*. Anterior is to the left. A γ of 0.5 was applied to the *nanos* channel for visualisation purposes. Scale bar = 50μm and applies to all panels. **(B)** Relative *nanos* and *pgc* signal enrichment (z-score) for the micrographs in (A) (gold), mean z-score lines and 95% confidence intervals for *y^1^ v^1^/w*; P{y^+^ v^+^ = (wt-osk-HA-bcd3’UTR)}attp40/+; P{w^+mC^=matalpha4-GAL-VP16}V37/+* (black line, positive control) and Oregon R (grey line, negative control). **(C)** Box plots of integrated anterior enrichment (integral of 15% anterior-most z-score values) for *nanos* (left) and pgc (right) in stage 1-2 embryos (right). Oregon R (negative control) in grey, median as grey dashed line. *y^1^ v^1^/w*; P{y^+^ v^+^ = (wt-osk-HA-bcd3’UTR)}attp40/+; P{w^+mC^=matalpha4-GAL-VP16}V37/+* (positive control) in black, median as solid black line. Significant difference from positive and negative control distributions (p-value (< 0.05) estimated using a bootstrap resampling test (10,000 iterations), comparing the observed difference in means between sample and control to a null distribution generated by resampling with replacement) indicated by asterisk and cross, respectively. Circles indicate interpretation of data: mRNA integrated anterior enrichment in variant significantly higher than negative control (dark golden), between positive and negative control (light golden), or significantly lower than positive control (grey).

Deleting either the OSK or the LOTUS domain significantly decreased the anterior enrichment of both mRNAs (Fig. 4C). However, the ΔLOTUS variant, but not the ΔOSK variant, showed HA-*nanos* and HA-*pgc* colocalisation comparable to that of WT Oskar, as measured by the Pearson correlation between the HA and *nanos* and *pgc* signal intensities of the anterior pixels (Fig. S4). Interestingly, the monomeric cricket LOTUS substitution failed, but the dimeric HlyU and LisH domain substitutions succeeded, in rescuing the mRNA recruitment defect (Fig. 4C). All substitutions, nonetheless, showed significantly lower degrees of recruitment of both studied mRNAs than controls (Fig. S4). None of the LOTUS variants were able to enrich *nanos* mRNA to wild type levels (Fig. 4C), and none of the LOTUS variants were able to enrich *pgc* mRNA to wild type levels (Fig. 4C) except for the HlyU substitution, which was capable of dimerisation *in vitro* (Fig. S1D). Despite their inability to sufficiently recruit *nanos* mRNA to the germ plasm, five of the LOTUS variants (DLOTUS, S158R, T196S, R215E, and the triple mutant) nevertheless showed apparently normal colocalisation of Oskar and *nanos* mRNA based on the signal intensity Pearson correlation (Fig. S4). Similarly, three of the LOTUS variants unable to recruit sufficient *pgc* mRNA to the germ plasm (DLOTUS, T195S and R215E) showed apparently normal colocalisation of Oskar and *pgc* mRNA (Fig. S4). These results suggest that the LOTUS domain is involved in mRNA recruitment, with dimerisation potentially being at least one aspect of its molecular mechanism.

## DISCUSSION

### Ectopic and endogenous Oskar may compete to localize germ plasm components

In *D. melanogaster*, formation of primordial germ cells, called pole cells, occurs in early embryogenesis and requires the products of multiple genes^8,54–57^. These genes act in a stepwise manner to enrich posterior accumulation of germ plasm components: Of these, only *oskar* is sufficient to ectopically trigger assembly of functional germ plasm ^10,12^. Wild-type anteriorly targeted *oskar* induced anterior pole cells in 15.1% of embryos (Fig. 2B; Table S2). Interestingly, *oskar*-*bcd*3’UTR embryos formed an average of 20.5 anterior pole cells and 24.3 posterior pole cells, each of which is about half of the average of 39.6 posterior pole cells seen in wild-type (Oregon R) embryos (Fig. 2C). This suggests that there may be competition between the ectopically targeted and the endogenous Oskar proteins for limiting germ plasm components. In some instances, we observed a phenotype that we scored as “anterior pole cell attempts” (Fig. 2A- B, Fig. S2). This phenotype is reminiscent of pole bud initiation, but not completion, observed at the posterior of *germ cell-less* (*gcl*) knockdown embryos ^58^, at the anterior of *gcl-bcd*3’UTR embryos ^59^, and in *osk*^301^, a temperature-sensitive *oskar* mutant ^37^. We speculate that these pole cell attempts occur in cases in which there is enough localised germ plasm to promote the initiation of pole cell cellularisation, but not enough to ensure completion of cellularisation nor complete adoption of germ cell fate.

### The relevance of Oskar dimerisation to pole cell formation

One of the strongest *in vitro* predictions regarding the molecular mechanism of Oskar is that the LOTUS domain can dimerised^21,22^. LOTUS dimerisation was interpreted by Jeske and colleagues ^21^to be mediated by residues D197, L200, and R215, in addition to the β sheets of the domain (residues 205-210, 215-219) ^21^. Congruently, dimerisation was reported by Yang et al ^22^ to be mediated by a hydrophobic patch including L200 and A207, and by polar interactions between D197 and R215. Our previous bioinformatic analyses predicted that the properties of these residues (electrostatic for D197 and R215; hydrophobic for L200) were highly conserved across insect Oskar proteins whose LOTUS domain is predicted to dimerised^60^. The dimer was disrupted *in vitro* by mutations S210P ^22^ and R215E ^21^, both of which were assessed *in vivo* herein. Our *in vivo* analysis includes two substitutions that preserve some of the original amino acid properties (i.e. L165A and T196S), which indeed did not disrupt LOTUS domain dimerisation in a bacterial two-hybrid assay (Fig. S1D). The remaining substitutions change the water affinity or charge of the targeted amino acid. Though predicted to be disruptive, D155A and S158R did not affect LOTUS dimerisation *in vitro* (Fig. S1D). More surprisingly, the triple mutant T196E-D197V-R215I could dimerise *in vitro*, even though the shielded hydrophobic T196 is converted into a charged residue that would be predicted to repel its counterpart on the participating LOTUS monomer, and the polar interactions between D197 and R215 are abolished (Fig. S1D). While flipping the charge in R215E ablated dimerisation ^21^ (Fig. S1D), turning the arginine into an uncharged glutamine in R215Q allowed dimerisation at the permissive temperature (RT). The R215Q substitution might, therefore, weaken dimerisation by abolishing the polar interaction with D197, but is not disruptive under permissive conditions.

Deletion of either the OSK or the LOTUS domain severely affected the ability of *oskar* to induce anterior pole cells *in vivo* (Fig. 2B). We observed no anterior pole cells in ΔOSK. However, even variants with an intact OSK domain displayed few or no anterior pole cells (Fig. 2B). Therefore, the OSK domain seems necessary but not sufficient for ectopic pole cell formation. The LOTUS domain seems to be important, but not strictly necessary for pole cell formation, since ΔLOTUS showed a low incidence of pole cells (Fig. 2B), albeit significantly lower than that of wild-type Oskar (Fig. 2B). LOTUS domain substitutions with the exogenous dimerisation domains HlyU from *Vibrio cholera* and LisH domain from *D. melanogaster* fared even worse than ΔLOTUS (Fig. 2B), indicating that dimerisation is not the sole function of the LOTUS domain in pole cell formation. Interestingly, the L165A and T196S mutations, which do not greatly affect the properties of the original amino acids, nonetheless showed significantly lower anterior pole cell incidence than controls (Fig. 2B), suggesting that even subtle changes to the LOTUS domain can have a large effect on the ability of Oskar to induce pole cells. Overall, we conclude that ectopic pole cell formation is an inefficient, exacting process that can be disrupted even by minor perturbations. Both well-folded Oskar domains appear to work synergistically to achieve pole cell formation, but only OSK seems to be strictly required.

### Robustness of the Oskar-Vasa interaction in vivo

Genetic evidence supports the hypothesis that Vasa’s initial germ plasm recruitment relies on interaction with Oskar^10,61–64^. Numerous *in vitro* assays corroborate a direct *in vitro* interaction between Oskar and Vasa ^21,36,39–41^. Nearly 30 years ago, the Oskar-Vasa interaction was attributed to the C-terminal fragment of Oskar (residues 290-606), which contains the OSK domain (residues 401-606), based on a yeast two-hybrid assay^39^. In contrast, a strong *in vitro* LOTUS domain-Vasa interaction was subsequently supported by the results of GST-pulldowns ^21,36,52^, isothermal titration calorimetry, yeast two-hybrid assays, and a co-crystal of LOTUS in complex with Vasa ^21,36^. Another *in vitro* study mapped the Oskar-Vasa interaction to a region of Vasa distinct from that predicted to bind the Oskar LOTUS domain ^38^. These *in vitro* data can be interpreted as predicting the existence of multiple Oskar-Vasa binding interfaces, involving both folded Oskar domains and various Vasa regions. In a heterologous *in vivo* system of *Drosophila* cultured cells, full-length Oskar recruited Vasa from the cytoplasm to Oskar-positive nuclear speckles ^36,65^. Deletion of the Oskar LOTUS domain diminished, but did not completely abolish, Vasa recruitment to nuclear speckles^36^, raising the possibility of a weak, LOTUS domain-independent Oskar-Vasa interaction *in vivo*. Our *in vivo* assessment of ectopic Vasa recruitment shows that neither the LOTUS domain nor the OSK domain are absolutely required for anterior Vasa enrichment, although both ΔLOTUS and ΔOSK display a “delayed enrichment” phenotype: worse than controls in S10 oocytes but reaching control levels by embryonic stage 1-2 (Fig. 3C).This could correspond to reduced recruitment efficiency.

In accordance with the observed dispensability of the LOTUS domain for ectopic Vasa recruitment, point mutations in the LOTUS domain also do not substantially affect anterior Vasa enrichment (Fig. 3C). The reported co-crystal structure predicted that the LOTUS domain-Vasa binding interface involves the pocket formed by the α2 and α5 helices ^36^. The interface 2 point mutants included in our analysis, D155, S158, and L165 were predicted to contact Vasa, but were not tested directly in previous studies ^36^. Indeed, substitutions D155A, S158R, and even the hydrophobic-to-hydrophobic L165A abrogated Oskar-Vasa interaction *in vitro* in our bacterial two-hybrid assay (Fig. S1D). However, *in vivo*, only the triple mutant T196E-D197V-R215I failed to localise Vasa to the anterior (Fig. 3C). This triple mutant phenotype is not easily interpretable as directly disrupting *in vivo* Oskar-Vasa interactions, given that the specific perturbations are far from the predicted Vasa-binding site and, additionally, do not affect Oskar-Vasa interaction or Oskar homodimerisation *in vitro* (Fig. S1D). The D155A, S158R, L165A, D197N, and R215E substitutions, which affect Oskar-Vasa interaction *in vitro* (Fig. S1D), are all able to recruit Vasa to the anterior *in vivo* (Fig. 3C). However, D155A, D197N, S210P, and R215Q show a “defective retention” phenotype (better Vasa enrichment in S10 oocytes than in S1-2 embryos; Fig. 3C), which might be caused directly or indirectly by sub-optimal Oskar-Vasa interactions. Vasa residue F504, predicted to contact the Oskar LOTUS domain based on the co-crystal structure, was previously shown to be important for *in vitro* Vasa-LOTUS domain interaction ^36^. This residue is also important for *in vivo* Vasa localisation to nurse cell nuage and to oocyte germ plasm^36^. Vasa mutants carrying mutations in this residue failed to rescue the hatching defect of a *vasa* mutant ^36^. The role of the reciprocal mutations on the Oskar LOTUS domain were, however, not previously tested *in vivo*, leaving open the possibility that the effect of Vasa mutations at F504 and F508 *in vivo* may be mediated by a mechanism other than binding to the Oskar LOTUS domain.

Overall, our results suggest a discrepancy between the *in vitro* and *in vivo* results on the interactions between Oskar and Vasa proteins, which highlights the complexity of the intracellular environment. Additionally, we suggest that Vasa recruitment to the germ plasm is robust, with redundant mechanisms of recruitment and retention. We favour a model where, in addition to the strong LOTUS domain-Vasa interaction, one or more alternative, perhaps weaker, Oskar-Vasa binding sites can compensate for the loss or disruption of the LOTUS domain-Vasa binding domain. If, alternatively, there is an indirect association between Oskar and Vasa, there is evidence that such an association would still be mediated via protein-protein interactions, since Vasa variants that have reduced RNA-binding activity are nonetheless able to localise to the germ plasm^61^.

### The LOTUS domain mediates nanos and pgc mRNA recruitment to the germ plasm

In addition to directly interacting with proteins, Oskar is an RNA-binding molecule ^21^. Bioinformatic analysis predicts that the LOTUS domain of Oskar would have RNA binding ability^17,20,60^. The OSK domain, on the other hand, which resembles a catalytically inactive SGNH hydrolase ^21,22^, does not have obvious predicted RNA binding capacity based on primary or secondary structure. These predictions notwithstanding, an *in vitro* mRNA pull down showed co-elution of the OSK domain, but not of the LOTUS domain, with mRNAs ^21^, and a nitrocellulose- filter-binding assay showed that the OSK domain, but not the LOTUS domain, was able to bind a nonspecific RNA oligo ^21^. Further, Yang and colleagues^22^ found that the OSK domain, but not the LOTUS domain, specifically recognised and bound the *oskar* and *nanos* 3’UTR ^22^ (but not the *pgc* 3’UTR^22^) in an electrophoretic mobility shift assay. Its crystal structure suggests that most of the surface of the OSK domain is positively charged or hydrophobic, which is consistent with RNA binding abilities ^21^. In agreement with these results, we found that ΔOSK fails to recruit *nanos* or *pgc* mRNAs to the anterior of embryos (Fig. 4C), consistent with previous observations^66^.

More interesting, however, was our observation that the ΔLOTUS Oskar variant also impaired mRNA localisation (Fig. 4C). This effect could be mediated by direct LOTUS domain- mRNA binding, by LOTUS domain dimerisation, or by an alternative, indirect method of mRNA recruitment. In support of the first hypothesis, the Oskar LOTUS domain was shown to interact with RNA G-quadruplex (G4) tertiary structures with high affinity by ELISA ^52^. ELISA showed that G4 RNA is specifically recognised by the LOTUS domains of both *D. melanogaster* Oskar and mouse TDRD5, as well as additional LOTUS domains from proteins in mammals, arthropods, plants, and bacteria ^52^, suggesting the interaction is ancient and widespread ^52^. We note that Jeske and colleagues ^21^ previously used UV-crosslinking to show that Oskar protein immunoprecipitated from embryo lysates was associated with *nanos*, *pgc*, and *gcl* mRNAs ^21^, an interaction that could in principle have been contributed to or mediated by the LOTUS domain. Hence, we hypothesise that both OSK and LOTUS are involved in mRNA recruitment *in vivo*. Consistent with a role for dimerisation of the LOTUS domain in nucleic acid recruitment, mutations in both LOTUS domain putative dimerisation interfaces disrupted recruitment of *nanos* and *pgc* mRNAs (Fig. 4C). LOTUS substitution with the monomeric Gb LOTUS impaired mRNA anterior enrichment, while substitution with the dimeric Vc HlyU partially rescued *nanos* and fully rescued *pgc* localisation (Fig. 4C). Substitution with the dimeric Dm LisH partially rescued recruitment of both mRNAs (Fig. 4C). These results suggest that dimerisation at the LOTUS domain, while not solely responsible for *nanos* and *pgc* interaction with Oskar, constitutes at least one aspect of the mechanism of Oskar-mediated *nanos* and *pgc* recruitment to the germ plasm. Consistent with this interpretation, the monomeric S210P LOTUS mutant did not show *oskar* 3’UTR RNA-binding activity *in vitro* ^22^.

While our results suggest that Oskar dimerisation via the LOTUS domain plays a role in mRNA recruitment, the solved structures of the LOTUS domain ^21,22^ suggest that its α3 helix, the surface that interacts with nucleic acids in the similarly folded winged-helix domain transcription factor MecI ^67^, would be disrupted by LOTUS dimerisation, inconsistent with a model of simultaneous dimerisation and direct mRNA binding. At the same time, the UV crosslinking and mRNA pull-down assay that identified co-elution of Oskar with mRNAs did not show co-purification of Vasa with Oskar ^21^. The absence of Vasa in the eluates in that study was attributed to the stringent purification conditions and low efficiency UV crosslinking ^21^, but an alternative explanation could be that Oskar is not often simultaneously bound to Vasa and mRNAs via its LOTUS domain, consistent with a model of competition between Vasa, mRNAs, and other Oskar molecules for the same LOTUS sites. We therefore propose a hypothesis that is consistent both with previously reported *in vitro* observations and with our *in vivo* investigation (Fig. 4C): that both interface 1 and interface 2 have the potential to bind mRNAs, as well as distinct functions in homodimerisation and Vasa binding.

### A non-modular model of Oskar molecular mechanism

Current models posit that each well-folded Oskar domain has separable molecular functions, namely that OSK performs mRNA binding and LOTUS enables protein-protein interactions. Our *in vivo* characterisation of a series of transgenic variants inspired by *in vitro* structural work is not consistent with a strictly modular view of Oskar’s function. We show instead that the entire LOTUS domain is dispensable for ectopic Vasa recruitment, which suggests a robust, redundant mechanism for Vasa germ plasm enrichment, potentially involving OSK and the interdomain. Additionally, we present strong *in vivo* evidence that LOTUS is involved in the recruitment of *nanos* and *pgc* mRNAs. This could point to a previously overlooked *in vivo* RNA-binding ability of the LOTUS domain and is also consistent with a synergistic mechanism for mRNA accumulation, involving multiple Oskar domains.

Oskar’s non-modular molecular mechanism is also consistent with its phase-separating properties ^65^. Oskar’s interdomain contains an 160 amino acid-long intrinsically disordered region and the first 47 amino acids of the LOTUS domain contain a low-complexity region, which were originally speculated to drive the observed condensation of Oskar into phase-separating granules^65^. In cultured *Drosophila* cells ^65^, no single domain (LOTUS, OSK, intrinsically disordered region, or low-complexity) was responsible for condensation of Oskar into liquid-like droplets ^65^. Deletion of any of these domains, however, reduced the efficiency of condensation ^65^, suggesting a redundant and synergistic function of Oskar’s domains in mediating condensation. We favour a non-modular view of Oskar’s function beyond phase-separation. We provide evidence that folded domains of Osk are neither necessary nor sufficient for their well-characterised germ plasm assembly functions *in vivo*, suggesting that redundant (in the case of Vasa) or synergistic (in the case of *nanos* and *pgc*) molecular mechanisms govern germ plasm recruitment. We speculate that such functional redundancy and synergy might have evolved to safeguard the critical process of establishing a viable germ line.

## RESOURCE AVAILABILITY

<u>Lead contact</u>: Further information and requests for resources and reagents should be directed to and will be fulfilled by the lead contact, Cassandra G. Extavour.

Materials availability: All materials generated and reported here are available from the lead contact upon request.

<u>Data and code availability</u>: All original code has been deposited at github at https://github.com/Anastasiarep/LOTUS_mutant_analysis.git (commit ID 0de8b5a) and is publicly available as of the date of publication. Any additional information required to reanalyze the data reported in this paper is available from the lead contact upon request.

## ACKNOWLEDGMENTS

We thank Rachelle Gaudet, Victoria D’Souza, Chandrashekar Kuyyamudi, Tarun Kumar, and Suhrid Ghosh for helpful discussions. A.R. was supported by the Boehringer Ingelheim Fonds Ph.D. fellowship. E.L.R. was supported by a Herchel Smith Graduate Fellowship, the Harvard Quantitative Biology Initiative, and the NSF-Simons Center for Mathematical and Statistical Analysis of Biology at Harvard (NSF award number 1764269). This work was supported by NIH R01 award 5R01GM143611-02 to C.G.E., who is a Howard Hughes Medical Institute investigator.

## AUTHOR CONTRIBUTIONS

**Anastasia Repouliou** performed all *in* vivo investigations, acquired and curated microscopy data, conceptualised, developed, and executed computational, analytical, and statistical analyses, generated figures and data visualisation, and prepared and edited the manuscript. **John R. Srouji** conceptualised the project, carried out all structural and biochemical investigations, developed relevant biochemical methodologies, generated all *Drosophila melanogaster* transgenic lines, and prepared an initial manuscript draft. **Emily L. Rivard** generated and validated an antibody reagent. **Andrés E. Leschziner** conceptualised the project, secured funding, provided resources, supervised and advised JRS. **Cassandra G. Extavour** conceptualised the project, secured funding, provided resources, administered the project, supervised and advised AR, JRS, and ELR, and reviewed and edited the manuscript. All authors have reviewed and approved the final manuscript.

## SUPPLEMENTAL INFORMATION

Document S1: contains Figure S1 – S5, Tables S1 – S5, and Key Resources Table.

## MATERIALS & METHODS

### Cloning related to Oskar LOTUS domains

Initial sequence boundaries for the Oskar LOTUS domain were set based on sequence similarity. A multiple sequence alignment of Drosophilid and cricket Oskar protein homologs was generated with MAFFT ^70^using the L-INS-i setting, default parameters, and Oskar protein sequences from the following species: *D. melanogaster* (UniProt Acc. No. P25158), *D. sechellia* (UniProt Acc. No. B4HKZ1), *D. simulans* (UniProt Acc. No. B4QXC8), *D. yakuba* (UniProt Acc. No. B4PTX6), *D. immigrans* (UniProt Acc. No. A1Y1T7), *D. virilis* (UniProt Acc. No. Q24741), *D. pseudoobscura* (UniProt Acc. No. Q295Q4), *D. erecta* (UniProt Acc. No. B3P1W4), *D. ananassae* (UniProt Acc. No. B3LZ06), *D. persimilis* (UniProt Acc. No. B4GFV0), *D. willistoni* (UniProt Acc. No. B4N815), *D. mojavensis* (UniProt Acc. No. B4K9E4), *D. grimshawi* (UniProt Acc. No. B4JTJ1), and *Gryllus bimaculatus* (UniProt Acc. No. K4MTL4). Based on the alignment, *D. melanogaster* Oskar residues 139 to 241 (abbreviated as Losk139-241) and *G. bimaculatus* Oskar residues 1 to 90 (abbreviated as Gbosk1-90) were designated as the beginning and end, respectively, of the LOTUS domain, because sequence similarity rapidly deteriorated beyond these points.

Almost all bacterial expression constructs were derived from pET-43.1b(+) (Novagen) and modified via circular polymerase extension cloning (CPEC) ^71,72^. The N-terminal NusA solubility partner, polyhistidine tag, and TEV cleavage site elements were removed and Losk139-241 followed by a 3C protease cleavage site (LEVLFQ|GP) and a decahistidine purification tag was introduced, yielding pET-Losk139-241_C10H. This expression vector subsequently served as the DNA template when introducing LOTUS domain point and deletion mutations via site-directed mutagenesis. To generate a bacterial expression construct for the cricket LOTUS domain, a published reagent (pET-151-Gbosk1-440^50^) was modified to encode Gbosk1-90 downstream of a hexahistidine tag and a TEV protease cleavage site, yielding pET-151-Gbosk1-90.

Transgenic inserts encoding anteriorly localised *D. melanogaster oskar* were based on a previously published reagent ^73^ that contains a fusion of the *D. melanogaster oskar* protein-coding sequence with the *D. melanogaster bicoid* 3’-UTR. This fusion was cloned and inserted into the *D. melanogaster* transgenic vector pVALIUM22 ^48,74,75^, yielding pVAL22-Dmosk-bcd3UTR ^50^. A C-terminal hemagglutinin (HA) epitope tag (YPYDVPDA) was inserted, yielding pVAL22- Dmosk-HA-bcd3UTR. pVAL22-Dmosk-HA-bcd3UTR served as the template for subsequent transgenic constructs encoding deletions of the LOTUS (residues 140 to 241) or OSK (residues 401 to 606) domains, yielding pVAL22-Dmosk-ΔLOTUS-HA-bcd3UTR and pVAL22-Dmosk-Δlipase-HA- bcd3UTR, respectively. To replace the LOTUS domain with exogenous protein sequences, the necessary DNA fragments were inserted into pVAL22-Dmosk-ΔLOTUS-HA-bcd3UTR via isothermal ligation assembly ^76^. The *Vibrio cholera* transcription factor HlyU (residues 1 to 109 from UniProt Acc. No. C3LST3; abbreviated as VcHlyU1-109) was cloned from a plasmid purchased from the DNASU Plasmid Repository (Arizona State University). The LisH domain from *D. melanogaster lis1* (residues 1 to 86 from UniProt Acc. No. Q7KNS3; abbreviated as DmLisH1-86) was amplified from a wild type *D. melanogaster* (Oregon R) ovarian cDNA library. The *G. bimaculatus oskar* LOTUS domain (residues 1 to 90 from UniProt Acc. No. K4MTL4) was cloned from pET-151-Gbosk1-440 ^50^.

### Expression and purification of Oskar LOTUS domains

BL21(DE3) *E. coli* were transformed with pET-Losk139-241_C10H and a single colony was used to inoculate 50 mL Luria-Bertani (LB) medium containing 100 µg/mL ampicillin for overnight growth at 37°C and 250 rpm shaking in a 250-mL flask. The following day, two 2-L LB/ampicillin cultures were each inoculated with 20 mL of the overnight starter culture and grown at 37°C with 225 rpm shaking until reaching OD600 ∼0.6. The culture was then transferred to 20 – 22°C (room temperature) and isopropyl-β-D-thiogalactopyranoside (IPTG) added to a final concentration of 0.5 mM for overnight expression with 225 rpm shaking. Cells were harvested from all 4 L of culture via centrifugation at 4200 rpm, 4°C in a JS-4.2 rotor (Beckman Coulter) for 25 minutes.

Harvested cells were resuspended with lysis buffer (50 mM Tris-HCl pH 8.0, 300 mM NaCl, 20 mM imidazole pH 8.0, 10% (v/v) glycerol, 10 mM 2-mercaptoethanol, 1 µg/mL pepstatin-A, 1 µg/µL aprotinin, 1 mM PMSF, 0.4 mg/mL hen egg white lysozyme) at a ratio of 5 mL buffer to 1 g of cells. Cells were lysed via sonication and the resulting crude lysate centrifuged at 20,000 rpm, 4°C for 45 minutes in a JA-20 rotor (Beckman Coulter). The resulting supernatant (clarified lysate) was applied to 5 mL Ni-NTA agarose (QIAGEN or Thermo Scientific) and agitated gently at 4°C for 1 hour to allow for batch affinity binding. Then, the slurry was passed through a column support to collect and separate the Ni-NTA agarose resin from the flowthrough. The resin was washed with 20 column volumes of lysis buffer at ∼1 mL/min by gravity. Losk139- 241_C10H was eluted in four separate 10-mL fractions with elution buffer (50 mM Tris pH 8.0, 300 mM NaCl, 500 mM imidazole pH 8.0, 10% (v/v) glycerol, 10 mM 2-mercaptoethanol) and its purity assessed by SDS-PAGE.

The polyhistidine purification tag was removed via overnight treatment of Losk139- 241_C10H with 1:100 m:m glutathione-S-transferase tagged 3C protease at 4°C, resulting in Losk139-241 (note that this leaves a remnant cleavage site sequence of LEVLFQ at the C- terminus). Subsequent purification of Losk139-241 via gel filtration (HiLoad Superdex 75 16/60 pg (GE Healthcare) using 20 mM Tris pH 8.0, 200 mM NaCl, 5 mM 2-mercaptoethanol) yields a Gaussian distribution of pure protein centered at the retention volume of ∼65 mL. The final purity of Losk139-241 was determined via SDS-PAGE and the protein concentrated with an Amicon Ultracel 10K MWCO regenerated cellulose device (Millipore).

To express N-terminally polyhistidine tagged Gbosk1-90 protein, BL21(DE3) *E. coli* were transformed with pET-151-Gbosk1-90 and protein production induced overnight at room temperature as described above for Losk139-241_C10H. The expressing cells were lysed and Gbosk1-90 was purified on Ni-NTA as described for Losk139-241_C10H. Afterwards, the polyhistidine affinity tag was removed via overnight TEV protease 113 cleavage (1:100 m:m protease:substrate) at 4°C.

To subsequently remove polyhistidine-tagged TEV protease and uncleaved substrate, the overnight TEV digestion was repeatedly diluted and concentrated with Ni-NTA lysis buffer (thus reducing the imidazole concentration) and passed through a Ni-NTA agarose column. The flowthrough from this second Ni-NTA purification was subsequently concentrated and purified via size exclusion chromatography on a HiLoad Superdex 75 16/60 pg column (GE Healthcare) with 20 mM Tris pH 8.0, 200 mM NaCl, 5 mM 2-mercaptoethanol. Gbosk1-90 elutes at a retention volume of ∼77 mL. The final purity of Gbosk1-90 was determined via SDS-PAGE and the protein concentrated with an Amicon Ultracel 3K MWCO regenerated cellulose device (Millipore).

*Crystallisation, data collection, and structure solution for Losk139-241 crystal forms 1-4* Crystals of Losk139-241 belonging to form 1 appeared after approximately three weeks via either sitting or hanging drop vapor diffusion at 4°C. Purified Losk139-241 (5 mg/mL) and reservoir solution (0.1 M MgCl2, 50 mM Tris pH 8.5 or 9.0, 17 or 18% (w/v) PEG 3350) were combined at a volumetric ratio of 1:1 or 1:2 protein:reservoir. For sitting drop vapor diffusion, crystals grew in a final drop volume of 1.5 µL exposed to a reservoir volume of 70 µL. For hanging drop vapor diffusion, crystals grew in a final drop volume spanning 2-6 µL exposed to a reservoir volume of 500 µL. To cryoprotect these crystals before plunge-freezing into liquid nitrogen, they were transferred from the crystallisation drop with a cryo-loop into five consecutive 5-µL drops composed of reservoir solution stepwise-diluted with cryoprotectant (0.1 M MgCl2, 50 mM Tris pH 8.5 or 9.0, 20% (w/v) PEG 3350, 25% (v/v) glycerol). Each crystal spent ∼1 minute in each drop before freezing.

Crystal forms 2 and 3 of Losk139-241 were discovered while inspecting three-year-old commercial sitting drop screens stored at 4 °C. Purified Losk139-241 (0.5 µL of 10 mg/mL) and reservoir solution (0.5 µL) were combined in a total drop exposed to a reservoir volume of 70 µL. Crystal form 2 grew from a reservoir solution of 0.05 M sodium sulfate, 0.05 M lithium sulfate, 0.05 M Tris pH 8.5, 30% (w/v) PEG 400. Crystal form 3 grew from a reservoir solution of 0.2 M sodium acetate, 0.1 M Tris pH 8.5, 16% (w/v) PEG 4000. Crystal forms 2 and 3 could not be reproduced in a 3-month period under the above-described conditions. Form 2 crystals were plunge frozen in liquid nitrogen directly from the drop using a cryo-loop. Form 3 crystals were quickly soaked (less than 10 seconds) in reservoir solution supplemented with 25% (v/v) glycerol before freezing in liquid nitrogen.

To derivatise the LOTUS domain for intended single anomalous dispersion (SAD) experiments, freshly prepared 1 mM mercury chloride (dissolved in MilliQ water, Millipore) was combined with 5.0 mg/mL purified Losk139-241 in a 5:1 molar ratio and incubated on ice for 25 minutes. The derivatisation reaction was then centrifuged at 20,800 x g and 4°C for 30 minutes to pellet any precipitated protein. To produce crystal form 4 via sitting drop diffusion, 1 µL of this Losk139-241 + HgCl2 sample was combined with 0.5 µL of reservoir solution (100 mM Tris pH 7.0, 150 mM KBr, 25-30% (w/v) PEG MME 2000) in a drop exposed to 70 µL of reservoir solution at 4 °C over 12 to 24 hours. These crystals were cryoprotected by quickly soaking (less than 10 seconds) in cryoprotectant (100 mM Tris pH 7.0, 150 mM KBr, 28% (w/v) PEG MME 2000, 20% (v/v) glycerol) before plunge-freezing in liquid nitrogen.

All X-ray diffraction data were collected at 100 Kelvin. Crystal form 1 data were collected with an ADSC Q315 detector and data for crystal forms 2-4 were collected with a Dectris Pilatus 6MF pixel array detector (ID24 beamline, Advanced Photon Source, Argonne, Illinois, U.S.A.). Data were processed in HKL2000 114; data statistics are shown in Supplementary Table 1. Initial phases for the Losk139-241 + HgCl2 structure (crystal form 4) were obtained via molecular replacement using MR-Rosetta ^77^ (as part of the PHENIX software package ^78,79^) and a search model template consisting of a monomeric homology model of Losk139-241 generated with MODELLER ^80^as part of the HHpred webserver ^81^. These phases were improved by incorporating SAD data collected at the mercury K-edge peak wavelength with MR-SAD and AutoSol as part of the PHENIX software package ^78,79^. This phased structure (crystal form 4) then served as the molecular replacement search model to phase native diffraction data collected from crystal forms 1-3 in Phaser ^82^. Any necessary model building was performed in COOT ^83^. Figures were generated with PyMOL^84^.

### Analytical size exclusion chromatography (analytical SEC) and SEC multiangle light scattering (SEC-MALS)

All analytical SEC experiments were conducted on the same day in rapid succession on an ÄKTA Micro system equipped with a Superose 6 PC 3.2/30 column (GE Healthcare). Fly and cricket Oskar LOTUS domains (Losk139-241 and Gbosk1-90, purified and concentrated as described above (“*Expression and purification of Oskar LOTUS domains*”) were each tested in triplicate at 5.0 mg/mL using identical run parameters that consisted of the following: a running buffer of 20 mM Tris pH 8.0 with either 200, 500, or 1000 mM NaCl and a flowrate of 0.05 mL/min. Gel filtration standards (BIO-RAD) were tested in all three buffers for subsequent determination of apparent molecular weight. Peak retention volumes were determined using UNICORN software (GE Healthcare).

A relationship exists between the extent to which a protein can penetrate a gel filtration matrix (the partition coefficient) and the logarithm of its molecular weight 91, 119. This partition coefficient (Kav) can be quantitatively expressed as follows: Kav = (Ve – Vo)/(Vt – Vo) where Ve is the elution (retention) volume of the sample protein, Vo is the column void volume (for a Superose 6 PC 3.2/30 column, this is 0.86 mL), and Vt is the total column volume (for a Superose 6 PC 3.2/30 column, this is 2.4 mL). The underlying assumption for this relationship to hold true is that the sample protein bears significant surface biochemical and overall geometric similarity to the standards. A calibration curve (for each running buffer) was generated in Python with a custom script using the determined retention peak volumes and known molecular weights for the gel filtration standards (GE Healthcare). This calibration curve was subsequently applied to Losk139- 241 and Gbosk1-90 analytical SEC data, yielding their apparent molecular weight.

For SEC-MALS analysis, fly and cricket Oskar LOTUS domains (Losk139-241 and Gbosk1-90, purified and concentrated as described above (“*Expression and purification of Oskar LOTUS domains*”) were tested at a concentration of 10 mg/mL using a running buffer consisting of 20 mM Tris pH 8.0, 200 mM NaCl. All samples were tested on an Agilent liquid chromatography system equipped with a Superdex 75 10/300 GL column (GE Healthcare) upstream of a DAWN HELEOS multiangle light scattering instrument with an accompanying Optilab T-rEX refractive index detector (Wyatt). SEC-MALS data were subsequently analysed using ASTRA 6.0.3.16 software and compared to a standard of bovine serum albumin.

### Bacterial two-hybrid analysis

All necessary bacterial two-hybrid strains (BTH101 and DHM1 *E. coli*) and plasmids (pKNT25, pUT18, pKT25, pUT18C, pKT25-zip, and pUT18C-zip) were purchased as components of the Bacterial Adenylate Cyclase-based Two-Hybrid (BACTH) kit (Euromedex). The two-hybrid analysis was conducted according to kit instructions. Fly short Oskar (Losk139-606) and fly Oskar LOTUS domain (Losk139-241) were inserted into pKNT25, pKT25, pUT18, and pUT18C via isothermal ligation assembly. The appropriate pair of plasmids (i.e. a plasmid encoding a T25 adenylate cyclase fragment and a plasmid encoding a T18 adenylate cyclase fragment) was transformed into both bacterial display strains (BTH101 and DHM1 *E. coli*) and the transformants plated on LB/agar supplemented with 100 µg/mL ampicillin, 50 µg/mL kanamycin, 40 µg/mL 5- bromo-4-chloro-3-indoyl-β-D-galactopyranoside (X-gal), and 0.5 mM isopropyl-β-D-thiogalactopyranoside (IPTG). These plates were incubated at either 30°C or room temperature (20–22°C). Colonies that subsequently appeared on the plates were scored for oligomerisation/interaction based on their color as compared to negative control colonies (*E. coli* transformed with empty pKNT25 and pUT18 vectors; these appear white) and positive control colonies (*E. coli* transformed with pKT25-zip and pUT18C-zip vectors; these appear blue). For later BACTH analysis of mutations in the fly Oskar LOTUS domain (Losk139-241), substitutions were only introduced into pKNT25-Losk139-241 and pUT18-Losk139-241 (prepared as described above) via RTW-PCR.

### Drosophila melanogaster stocks

The maternal driver line *w[*]; P{w[+mC]=matalpha4-GAL-VP16}V37* (stock number 7063) was obtained from the Bloomington Drosophila Stock Center (Indiana, U.S.A.). All transgenic constructs were injected by Genetic Services, Inc. (GSI, Cambridge, Massachusetts, U.S.A.) into *y[1] v[1]; attP40*. Additionally, GSI used *y[1] v[1]; attP40* when screening for transformants. *y[1] sc[*] v[1]; Sco/CyO* was a kind gift of the Perrimon lab (Harvard Medical School).

### Creation of chicken anti- D. melanogaster Vasa antibody

A polyclonal chicken anti-*D. melanogaster* Vasa antibody was generated using AbClonal (Woburn, MA, USA) custom antibodies services. AbClonal synthesized gene sequences of an antigen target consisting of part of the *D. melanogaster* Vasa protein (UniProt ID: M9PBB5, M1-E206) and cloned it into expression vectors. They purified these proteins and injected them into chickens for immunization. Total IgY was precipitated and purified from 10 egg yolks of immunized chickens.

Antibodies were verified in our laboratory with western blots using ovary protein lysates (generated from 30 pairs of ovaries squished in RIPA buffer with protein amount estimated using a Pierce Detergent Compatible Bradford Assay (ThermoFisher Scientific) (Fig. S5A, B). SDS-PAGE gels were run with lysates mixed and boiled with beta-mercaptoethanol and 5x Pierce Lane Marker Non-Reducing Sample Buffer (ThermoFisher Scientific). Precision Plus Dual Color (Bio-Rad, chemiluminescent blots) or Precision Plus Western C standards (Bio-Rad, fluorescent blots)were used as standards. Semi-dry transfer was conducted using a Bio-Rad Trans-Blot Turbo Transfer System (Bio-Rad) and the Trans-Blot Turbo RTA PVDF Transfer Kit (Bio-Rad).

For chemiluminescent blots, membranes were washed once for five minutes in 1xTBS-T (1xTBS + 0.1% Tween-20) and blocked for one hour in 1xTBST + 5% milk (Carnation Instant Nonfat Dry Milk). Membranes were washed twice for one minute, once for 15 minutes, and twice for five minutes in 1xTBST before incubation overnight at 4°C in 1:500 primary antibody in 1xTBST + 5% BSA + 0.03% sodium azide. The following day, membranes were washed three times for ten minutes in 1xTSBT, incubated in 1:5000 anti-chicken HRP secondary antibody (ThermoFisher Scientific) in 1xTBST + 5% milk at room temperature, and washed three times for ten minutes in TBST. The SuperSignal West Pico PLUS Chemiluminescent Substrate (ThermoFisher Scientific) was used for detection. For fluorescent blots, membranes were blocked once for ten minutes in Fluorescent Blot Blocking Buffer (Azure) and then incubated in 1:500 primary antibody for one hour at room temperature. Membranes were then washed two times for one minute and three times for five minutes in 10X Blot Washing Buffer (Azure) and incubated in 1:5000 anti-chicken 800 secondary antibody (Azure) in blocking buffer for one hour at room temperature. Following secondary antibody incubation, membranes were washed twice for one minute, three times for five minutes in washing buffer, and once more for five minutes in 1xPBS. Blots were dried before imaging using a CCD camera on a Sapphire Biomolecular Imager.

The antibody was additionally validated with immunostaining *vasa* null ovaries (*vas[PH165]/Df(2L)b87e25*) using the ovary staining protocol described below (see “*Antibody staining of oocytes*” (Fig. S5C).

### Antibody staining of oocytes

Homozygous transgenic lines containing *oskar* variants under UAS control were crossed to the maternal GAL4 driver line *w[*]; P{w[+mC]=matalpha4-GAL-VP16}V37*, yielding female progeny expressing the corresponding *oskar* transgene (henceforth, these progeny and their offspring are referred to by their associated transgene). These offspring were mated to each other and raised on standard fly medium supplemented with yeast at 25°C and 60% relative humidity for two or three days before their ovaries were dissected. Ovaries were dissected in 1X phosphate buffer saline (PBS), fixed in 4% (w/v) paraformaldehyde in 1X PBS for 20 minutes, washed in 1XPBS, washed twice for five minutes in PBST (1X PBS + 0.2% (v/v) Tween-20), permeabilised for one hour in PBSTT (PBST + 1% v/v TritonX-100), washed for five minutes in PBST, and blocked twice for one hour in 5% NGS in PBST, on a roller mixer at room temperature. A cocktail of primary antibodies (diluted in 5% NGS in PBST) was added to the ovaries for overnight staining at 4°C. The second day, the ovaries were washed and blocked sequentially for 10 minutes, 20 minutes, 30 minutes, and 40 minutes with 5% NGS in PBST at room temperature. Then, a cocktail of fluorescently labelled secondary antibodies and DAPI nuclear stain (all diluted in 5% NGS in PBST) was added for overnight staining at 4°C. The third day, after five 5-minute washes in PBST, completely stained ovaries were mounted in VECTASHIELD (Vector Laboratories) and imaged. Primary antibodies and working dilutions for stains were as follows: 1:50 mouse anti- hemagglutinin (HA) tag (AE008, ABclonal) and 1:1,000 rabbit anti-*D. melanogaster* Vasa (obtained from Dr. Prashanth Rangan (Icahn School of Medicine at Mount Sinai, New York, USA)). Secondary antibodies were used at a 1:500 dilution as follows: goat anti-mouse conjugated to Alexa 488 (Thermo Fisher Scientific, A-11001) and goat anti-rabbit conjugated to Alexa 647 (Thermo Fisher Scientific, A-21245). DNA was stained with DAPI (Sigma-Aldrich, D9542) at 1:1,000 dilution of a 10 mg/mL stock solution.

### Antibody staining of stage 1-2 embryos

Homozygous transgenic lines were crossed to the maternal driver line *w[*]; P{w[+mC]=matalpha4-GAL-VP16}V37*, yielding female progeny expressing the corresponding *oskar* transgene (henceforth, these F1 progeny and their F2 offspring are referred to by their associated transgene). These F1 progeny (males and females) were placed in a cage over an apple juice agar plate before their fertilised embryos (F2) were collected. For germ plasm accumulation visualisation at stage 1-2, embryos were collected after 1 hour of embryo laying. Embryos were rinsed with water into a mesh basket, dechorionated in fresh 100% (v/v) bleach for 2-3 minutes, and washed extensively with water afterwards. Embryos were then fixed in a 1:1 volume:volume mixture of heptane and 4% (w/v) paraformaldehyde at room temperature with nutation for 20 minutes. The bottom aqueous layer was removed, an equal volume of 100% methanol added, and the embryos devitellinised by vigorously shaking their container for 2-3 minutes. The top organic layer was removed, another volume of 100% methanol added, and the container vigorously shaken again. After letting the embryos sink, the entire liquid solution was removed and 100% methanol added; this was repeated two times and the fixed embryos stored at –20 °C in 100% methanol until staining. For staining, embryos were gradually rehydrated in 1X PBS, then washed, permeabilised, and blocked twice for 15 minutes in PBTB (1X PBS + 0.2% (v/v) TritonX-100 + 1 mg/mL bovine serum albumin (BSA)), on a roller. A cocktail of primary antibodies (diluted in 5% NGS in PBTB) was added to the embryos for overnight staining at 4°C. The second day, the embryos were washed four times for 15 minutes in PBTB at room temperature, then stained overnight with fluorescently labelled secondary antibodies and DAPI nuclear stain (all diluted in 5% NGS in PBTB) at 4°C. The third day, after four 15-minute washes in PBTB and a 15-minute wash in 1X PBS, completely stained embryos were mounted in Vectashield (Vector Laboratories) and imaged.

Primary antibodies and working dilutions for stains were as follows: 1:50 rat anti-hemagglutinin (HA) tag (Roche, 3F10) and 1:1,000 chicken anti-*D. melanogaster* Vasa (this paper: see “*Creation of chicken anti- D. melanogaster Vasa antibody”*). Secondary antibodies used at a 1:500 dilution were as follows: goat anti-rat conjugated to Alexa 488 (Thermo Fisher Scientific, A-11006) and goat anti-chicken conjugated to Alexa 633 (Thermo Fisher Scientific, A-21103). DNA was stained with DAPI (Sigma-Aldrich, D9542) at 1:1,000 dilution of a 10 mg/mL stock solution.

### Antibody staining of stage 5 embryos

Embryos were collected as described above in “*Antibody staining of stage 5 embryos from transgenic mothers”*. For pole cell visualisation at stage 5, embryos were collected after two hours of egg-laying followed by removal of the apple juice agar plate and incubation for three hours at 25°C for embryo aging. Embryos were collected as described above with one notable distinction: fresh 100% ethanol was used instead of methanol in the previously described protocol to allow easier visualisation of actin filaments. In the ethanol protocol, all steps were performed as in the methanol protocol, but the devitellinisation by vigorous shaking step was longer (ten minutes) to promote effective devitellinisation. Additionally, embryos prepared by the ethanol protocol were stained on the same day they were collected.

Primary antibodies and working dilutions were as follows: 1:50 mouse anti-hemagglutinin (HA) tag (AE008, ABclonal) and 1:1,000 rabbit anti-*D. melanogaster* Vasa (obtained from Dr. Prashanth Rangan (Icahn School of Medicine at Mount Sinai, New York, USA)). Secondary antibodies used at a 1:500 dilution were as follows: goat anti-mouse conjugated to Alexa 488 (Thermo Fisher Scientific, A-11001) and goat anti-rabbit conjugated to Alexa 647 (Thermo Fisher Scientific, A-21245). DNA was stained with DAPI (Sigma-Aldrich, D9542) at 1:1,000 dilution of a 10 mg/mL stock solution. Actin was stained with a 6.6 µM stock solution of Rhodamine Phalloidin (Thermo Fisher Scientific, R415) at 1:100 dilution.

### HCR™ RNA-FISH and imaging of stage 1-2 embryos from transgenic mothers

Embryos were collected as described for stage 1-2 antibody-stained embryos and stored in methanol. For combined antibody stain and FISH, a modified version of the HCR™ IF+HCR™ RNA-FISH protocol for sample in solution provided by Molecular Instruments was used. Embryos were gradually rehydrated in PBST (1X PBS + 0.2% (v/v) Tween-20), washed twice for five minutes in PBST, permeabilised for one hour in PBSTT (PBST + 1% v/v TritonX-100), washed for five minutes in PBST, and blocked twice for one hour in 5% NGS in PBST, on a roller mixer at room temperature. The mouse anti-HA primary antibody (diluted in 5% NGS in PBST) was added to the embryos for overnight staining at 4°C. The second day, the embryos were washed and blocked four times for 30 minutes with PBST at room temperature. The fluorescently labelled secondary antibody (diluted in 5% NGS in PBST) was added for three hours at room temperature with nutation. Then, the embryos were washed five times for five minutes in PBST, followed by a five-minute incubation in 5X SSCT (5X saline sodium citrate (SSC) buffer + 0.1% Tween-20), a ten-minute post-fixation with 500μL 4% formaldehyde in PBST, two washes in PBST, two washes in 5X SSCT, and a pre-hybridisation incubation of at least two hours with pre-warmed probe hybridisation buffer in a 37°C water bath. We observed that the longer the incubation, the better the yield of stained embryos (data not shown). After embryos had sunk to the bottom of the tube, all but 50μL of pre-hybridisation was removed and a 16 nM probe solution was added to the embryos. The embryos were incubated with the probes overnight in a 37°C water bath. The third day, the embryos were washed four times for five minutes with pre-warmed probe wash buffer in a 37°C water bath and washed once for five minutes with 5X SSCT at room temperature. Embryos were incubated for at least 30 minutes with amplification buffer at room temperature. Meanwhile, hairpins h1 and h2 corresponding to each probe were separately snap-cooled by heating at 95°C for 90 seconds and cooling to room temperature in a dark drawer for 30 minutes. After the pre-amplification incubation, the amplification buffer was removed and a 60 nM snap-cooled hairpin solution in 100μL amplification buffer was added to the embryos. The embryos were incubated in the dark at room temperature overnight. The fourth day, embryos were washed sequentially twice for five minutes, then twice for 30 minutes, then once for five minutes in 5X SSCT, incubated in a 1:1,000 dilution of a 10 mg/mL stock solution of DAPI in PBST for at least 30 minutes, washed five times for five minutes with 5X SSCT, mounted in Vectashield (Vector Laboratories) and imaged.

Primary antibody and working dilutions were as follows: 1:50 mouse anti-hemagglutinin (HA) tag (AE008, ABclonal). The secondary antibody used at a 1:500 dilution was goat anti-mouse conjugated to Alexa 488 (Thermo Fisher Scientific, A-11001). DNA was stained with DAPI (Sigma-Aldrich, D9542) at 1:1,000 dilution of a 10 mg/mL stock solution. Probes and concentrations for HCR™ RNA-FISH were 16 nM HCR™ RNA-FISH anti-*nanos* probe (Molecular Instruments) and 16 nM HCR™ RNA-FISH anti-*pgc* probe (Molecular Instruments). Hairpins and concentrations for HCR™ RNA-FISH were 60nM HCR™ RNA-FISH amplifier B5- 555 (Molecular Instruments) and 60nM HCR™ RNA-FISH amplifier B4-647 (Molecular Instruments).

### Confocal image acquisition and analysis

Confocal images were collected on a Nikon CSU-W1 spinning disc confocal microscope, using identical laser power and exposure time for each sample type. Only oocytes or embryos with detectable, localized HA signal above background were imaged and analysed. For each sample, a Z-stack was obtained. A custom FIJI macro script was used to generate a sum intensity projection, to rotate the image so that the anteroposterior axis was horizontal and the posterior was oriented to the right, and to manually trace the outline of each oocyte or embryo to generate a mask.

Then, a custom Python script was used to mask the image, calculate the z-score for each pixel (computed as the signal intensity of each pixel minus the mean intensity value for the whole mask, divided by the standard deviation of intensity values within the whole mask), and sum the z-score value for the anterior-most 15% pixels, defined as “anterior integrated enrichment”.

Additionally, out of the anterior-most 15% pixels, the pixels with the highest HA signal values for each micrograph were defined as follows: a threshold was set to correspond to the mean HA intensity value plus two standard deviations. The pixels with intensity values above that threshold were selected and turned into a mask. A Gaussian filter was applied to the image. The mask corresponding to the anterior pixels with the highest HA value was applied to the filtered image. If the resulting masked image contained at least two pixel values, the Pearson’s correlation coefficient for the HA signal and either the Vasa, the *nanos*, or *pgc* signal was calculated, measured as a proxy for signal colocalisation.

The HA and Vasa signal enrichment (z-score) across normalised anteroposterior axis plots were generated by extracting the anterior and posterior boundaries of each sample from the provided mask, z-signal values were interpolated to generate a standardised 300-point linear profile per channel, allowing consistent comparisons across samples of varying size and shape, and z-score profiles per genotype were aggregated along the normalised anteroposterior axis to plot the mean and 95% confidence intervals (±1.96 × standard error of the mean).

Samples of stage 5 embryos were visually inspected and scored as HA-positive, showing phenotypes of developmental defect, anterior pole cell attempt, or anterior pole cells. Anterior and posterior pole cells were manually counted.

To assess statistical significance, we performed a bootstrapped test using 10,000 iterations. For each iteration, data points were resampled with replacement from the original dataset to generate a null distribution of the test statistic (difference in means). The p-value was computed as the proportion of bootstrapped statistics equal to or more extreme than the observed value. A p-value below 0.05 was considered to be statistically significant (marked by an asterisk). All p-values reported here were obtained by bootstrapping, except for the anterior pole cell incidence p-values (Table S3) for which one-sided Fisher’s exact tests were used to assess whether the proportion of embryos showing anterior pole cells differed significantly between experimental conditions and control groups. 2x2 contingency tables were constructed using the number of embryos with and without the anterior pole cell phenotype in each group.

## REFERENCES

1. Geigy, R. (1931). Action de l’ultra-violet sur le pôle germinal dans l’oeuf de Drosophila melanogaster (Castration et mutabilité). Revue suisse de zoologie 38, 187–285.

2. Illmensee, K., and Mahowald, A.P. (1974). Transplantation of posterior polar plasm in Drosophila. Induction of germ cells at the anterior pole of the egg. Proc. Natl. Acad. Sci. 71, 1016–1020. 10.1073/pnas.71.4.1016.

3. Illmensee, K., Mahowald, A.P., and Loomis, M.R. (1976). The ontogeny of germ plasm during oogenesis in Drosophila. Dev Biol 49, 40–65. 10.1016/0012-1606(76)90257-8.

4. Illmensee, K., and Mahowald, A.P. (1976). The autonomous function of germ plasm in a somatic region of the Drosophila egg. Exp Cell Res 97, 127–140. 10.1016/0014-4827(76)90662-5.

5. Mahowald, A.P., Illmensee, K., and Turner, F.R. (1976). Interspecific transplantation of polar plasm between Drosophila embryos. J Cell Biology 70, 358–373. 10.1083/jcb.70.2.358.

6. Ewen-Campen, B., Schwager, E.E., and Extavour, C.G.M. (2010). The molecular machinery of germ line specification. Mol. Reprod. Dev. 77, 3–18. 10.1002/mrd.21091.

7. Houston, D.W., and King, M.L. (2000). Germ plasm and molecular determinants of germ cell fate. Curr. Top. Dev. Biol. 50, 155-IN2. 10.1016/s0070-2153(00)50008-8.

8. Lehmann, R., and Nüsslein-Volhard, C. (1986). Abdominal segmentation, pole cell formation, and embryonic polarity require the localized activity of oskar, a maternal gene in drosophila. Cell 47, 141–152. 10.1016/0092-8674(86)90375-2.

9. Markussen, F.H., Michon, A.M., Breitwieser, W., and Ephrussi, A. (1995). Translational control of oskar generates short OSK, the isoform that induces pole plasm assembly. Dev Camb Engl 121, 3723–3732.

10. Ephrussi, A., and Lehmann, R. (1992). Induction of germ cell formation by oskar. Nature 358, 387–392. 10.1038/358387a0.

11. Ephrussi, A., Dickinson, L.K., and Lehmann, R. (1991). oskar organizes the germ plasm and directs localization of the posterior determinant nanos. Cell 66, 37–50. 10.1016/0092-8674(91)90137-n.

12. Kim-Ha, J., Smith, J.L., and Macdonald, P.M. (1991). oskar mRNA is localized to the posterior pole of the Drosophila oocyte. Cell 66, 23–35. 10.1016/0092-8674(91)90136-m.

13. Smith, J.L., Wilson, J.E., and Macdonald, P.M. (1992). Overexpression of oskar directs ectopic activation of nanos and presumptive pole cell formation in Drosophila embryos. Cell 70, 849–859. 10.1016/0092-8674(92)90318-7.

14. Castagnetti, S., Hentze, M.W., Ephrussi, A., and Gebauer, F. (2000). Control of oskar mRNA translation by Bruno in a novel cell-free system from Drosophila ovaries. Development 127, 1063–1068.

15. Kim-Ha, J., Kerr, K., and Macdonald, P.M. (1995). Translational regulation of oskar mRNA by Bruno, an ovarian RNA-binding protein, is essential. Cell 81, 403–412. 10.1016/0092-8674(95)90393-3.

16. Gunkel, N., Yano, T., Markussen, F.-H., Olsen, L.C., and Ephrussi, A. (1998). Localization- dependent translation requires a functional interaction between the 5′ and 3′ ends of oskar mRNA. Gene Dev 12, 1652–1664. 10.1101/gad.12.11.1652.

17. Callebaut, I., and Mornon, J.-P. (2010). LOTUS, a new domain associated with small RNA pathways in the germline. Bioinformatics 26, 1140–1144. 10.1093/bioinformatics/btq122.

18. Vollmer, W., Joris, B., Charlier, P., and Foster, S. (2008). Bacterial peptidoglycan (murein) hydrolases. FEMS Microbiol. Rev. 32, 259–286. 10.1111/j.1574-6976.2007.00099.x.

19. Finn, R.D., Bateman, A., Clements, J., Coggill, P., Eberhardt, R.Y., Eddy, S.R., Heger, A., Hetherington, K., Holm, L., Mistry, J., et al. (2014). Pfam: the protein families database. Nucleic Acids Res. 42, D222–D230. 10.1093/nar/gkt1223.

20. Anantharaman, V., Zhang, D., and Aravind, L. (2010). OST-HTH: a novel predicted RNA- binding domain. Biol Direct 5, 13. 10.1186/1745-6150-5-13.

21. Jeske, M., Bordi, M., Glatt, S., Müller, S., Rybin, V., Müller, C.W., and Ephrussi, A. (2015). The crystal structure of the Drosophila germline inducer Oskar identifies two domains with distinct Vasa helicase- and RNA-binding activities. Cell Rep. 12, 587–598. 10.1016/j.celrep.2015.06.055.

22. Yang, N., Yu, Z., Hu, M., Wang, M., Lehmann, R., and Xu, R.-M. (2015). Structure of Drosophila Oskar reveals a novel RNA binding protein. Proc Natl Acad Sci U S A 112, 11541–11546. 10.1073/pnas.1515568112.

23. Aravind, L., Anantharaman, V., Balaji, S., Babu, M.M., and Iyer, L.M. (2005). The many faces of the helix-turn-helix domain: Transcription regulation and beyond*. FEMS Microbiol. Rev. 29, 231–262. 10.1016/j.fmrre.2004.12.008.

24. Gajiwala, K.S., and Burley, S.K. (2000). Winged helix proteins. Curr. Opin. Struct. Biol. 10, 110–116. 10.1016/s0959-440x(99)00057-3.

25. Harami, G.M., Gyimesi, M., and Kovács, M. (2013). From keys to bulldozers: expanding roles for winged helix domains in nucleic-acid-binding proteins. Trends Biochem. Sci. 38, 364–371. 10.1016/j.tibs.2013.04.006.

26. Mukherjee, D., Datta, A.B., and Chakrabarti, P. (2014). Crystal structure of HlyU, the hemolysin gene transcription activator, from Vibrio cholerae N16961 and functional implications. Biochim. Biophys. Acta (BBA) - Proteins Proteom. 1844, 2346–2354. 10.1016/j.bbapap.2014.09.020.

27. Williams, S.G., Attridge, S.R., and Manning, P.A. (1993). The transcriptional activator HIyU of Vibrio cholerae: nucleotide sequence and role in virulence gene expression. Mol. Microbiol. 9, 751–760. 10.1111/j.1365-2958.1993.tb01735.x.

28. Liu, Y., Manna, A., Li, R., Martin, W.E., Murphy, R.C., Cheung, A.L., and Zhang, G. (2001). Crystal structure of the SarR protein from Staphylococcus aureus. Proc. Natl. Acad. Sci. 98, 6877–6882. 10.1073/pnas.121013398.

29. Lachke, S.A., Alkuraya, F.S., Kneeland, S.C., Ohn, T., Aboukhalil, A., Howell, G.R., Saadi, I., Cavallesco, R., Yue, Y., Tsai, A.C.-H., et al. (2011). Mutations in the RNA Granule Component TDRD7 Cause Cataract and Glaucoma. Science 331, 1571–1576. 10.1126/science.1195970.

30. Cui, G., Botuyan, M.V., and Mer, G. (2013). 1H, 15N and 13C resonance assignments for the three LOTUS RNA binding domains of Tudor domain-containing protein TDRD7. Biomol. NMR Assign. 7, 79–83. 10.1007/s12104-012-9382-1.

31. Oppikofer, M., Kueng, S., Keusch, J.J., Hassler, M., Ladurner, A.G., Gut, H., and Gasser, S.M. (2013). Dimerization of Sir3 via its C-terminal winged helix domain is essential for yeast heterochromatin formation. EMBO J. 32, 437–449. 10.1038/emboj.2012.343.

32. Akoh, C.C., Lee, G.-C., Liaw, Y.-C., Huang, T.-H., and Shaw, J.-F. (2004). GDSL family of serine esterases/lipases. Prog. Lipid Res. 43, 534–552. 10.1016/j.plipres.2004.09.002.

33. Ašler, I.L., Ivić, N., Kovačić, F., Schell, S., Knorr, J., Krauss, U., Wilhelm, S., Kojić-Prodić, B., and Jaeger, K. (2010). Probing Enzyme Promiscuity of SGNH Hydrolases. ChemBioChem 11, 2158–2167. 10.1002/cbic.201000398.

34. Lo, Y.-C., Lin, S.-C., Shaw, J.-F., and Liaw, Y.-C. (2003). Crystal Structure of Escherichia coli Thioesterase I/Protease I/Lysophospholipase L1: Consensus Sequence Blocks Constitute the Catalytic Center of SGNH-hydrolases through a Conserved Hydrogen Bond Network. J. Mol. Biol. 330, 539–551. 10.1016/s0022-2836(03)00637-5.

35. Lo, Y.-C., Lin, S.-C., Shaw, J.-F., and Liaw, Y.-C. (2005). Substrate Specificities of Escherichia coli Thioesterase I/Protease I/Lysophospholipase L1 Are Governed by Its Switch Loop Movement †. Biochemistry 44, 1971–1979. 10.1021/bi048109x.

36. Jeske, M., Müller, C.W., and Ephrussi, A. (2017). The LOTUS domain is a conserved DEAD-box RNA helicase regulator essential for the recruitment of Vasa to the germ plasm and nuage. Genes Dev. 31, 939–952. 10.1101/gad.297051.117.

37. Hay, B., Jan, L.Y., and Jan, Y.N. (1990). Localization of vasa, a component of Drosophila polar granules, in maternal-effect mutants that alter embryonic anteroposterior polarity. Development 109, 425–433. 10.1242/dev.109.2.425.

38. Anne, J. (2010). Targeting and Anchoring Tudor in the Pole Plasm of the Drosophila Oocyte. Plos One 5, e14362. 10.1371/journal.pone.0014362.

39. Breitwieser, W., Markussen, F.H., Horstmann, H., and Ephrussi, A. (1996). Oskar protein interaction with Vasa represents an essential step in polar granule assembly. Genes Dev. 10, 2179–2188. 10.1101/gad.10.17.2179.

40. Carrera, P., Johnstone, O., Nakamura, A., Casanova, J., Jäckle, H., and Lasko, P. (2000). VASA Mediates Translation through Interaction with a Drosophila yIF2 Homolog. Mol. Cell 5, 181–187. 10.1016/s1097-2765(00)80414-1.

41. Hurd, T.R., Herrmann, B., Sauerwald, J., Sanny, J., Grosch, M., and Lehmann, R. (2016). Long Oskar Controls Mitochondrial Inheritance in Drosophila melanogaster. Dev. Cell 39, 560–571. 10.1016/j.devcel.2016.11.004.

42. Valdes, R., and Ackers, G.K. (1979). Study of Protein Subunit Association Equilibria by Elution Gel Chromatography. Methods Enzym. 61, 125–142. 10.1016/0076-6879(79)61011-x.

43. Wyatt, P.J. (1993). Light scattering and the absolute characterization of macromolecules. Anal. Chim. Acta 272, 1–40. 10.1016/0003-2670(93)80373-s.

44. Battesti, A., and Bouveret, E. (2012). The bacterial two-hybrid system based on adenylate cyclase reconstitution in Escherichia coli. Methods 58, 325–334. 10.1016/j.ymeth.2012.07.018.

45. Karimova, G., Pidoux, J., Ullmann, A., and Ladant, D. (1998). A bacterial two-hybrid system based on a reconstituted signal transduction pathway. Proc. Natl. Acad. Sci. 95, 5752–5756. 10.1073/pnas.95.10.5752.

46. Sahin, E., and Roberts, C.J. (2012). Therapeutic Proteins, Methods and Protocols. Methods Mol. Biol. (Clifton, NJ) 899, 403–423. 10.1007/978-1-61779-921-1_25.

47. Wang, C., and Lehmann, R. (1991). Nanos is the localized posterior determinant in Drosophila. Cell 66, 637–647. 10.1016/0092-8674(91)90110-k.

48. Markstein, M., Pitsouli, C., Villalta, C., Celniker, S.E., and Perrimon, N. (2008). Exploiting position effects and the gypsy retrovirus insulator to engineer precisely expressed transgenes. Nat Genet 40, 476–483. 10.1038/ng.101.

49. Brand, A.H., and Perrimon, N. (1993). Targeted gene expression as a means of altering cell fates and generating dominant phenotypes. Development 118, 401–415.

50. Ewen-Campen, B., Srouji, J.R., Schwager, E.E., and Extavour, C.G. (2012). oskar Predates the Evolution of Germ Plasm in Insects. Curr Biol 22, 2278–2283. 10.1016/j.cub.2012.10.019.

51. Kim, M.H., Cooper, D.R., Oleksy, A., Devedjiev, Y., Derewenda, U., Reiner, O., Otlewski, J., and Derewenda, Z.S. (2004). The Structure of the N-Terminal Domain of the Product of the Lissencephaly Gene Lis1 and Its Functional Implications. Structure 12, 987–998. 10.1016/j.str.2004.03.024.

52. Ding, D., Wei, C., Dong, K., Liu, J., Stanton, A., Xu, C., Min, J., Hu, J., and Chen, C. (2020). LOTUS domain is a novel class of G-rich and G-quadruplex RNA binding domain. Nucleic Acids Res. 48, 9262–9272. 10.1093/nar/gkaa652.

53. Millevoi, S., Moine, H., and Vagner, S. (2012). G-quadruplexes in RNA biology. Wiley Interdiscip. Rev.: RNA 3, 495–507. 10.1002/wrna.1113.

54. Boswell, R.E., and Mahowald, A.P. (1985). tudor, a gene required for assembly of the germ plasm in Drosophila melanogaster. Cell 43, 97–104. 10.1016/0092-8674(85)90015-7.

55. Boswell, R.E., Prout, M.E., and Steichen, J.C. (1991). Mutations in a newly identified Drosophila melanogaster gene, mago nashi, disrupt germ cell formation and result in the formation of mirror-image symmetrical double abdomen embryos. Development 113, 373–384. 10.1242/dev.113.1.373.

56. Manseau, L.J., and Schüpbach, T. (1989). cappuccino and spire: two unique maternal-effect loci required for both the anteroposterior and dorsoventral patterns of the Drosophila embryo. Genes Dev. 3, 1437–1452. 10.1101/gad.3.9.1437.

57. Schüpbach, T., and Wieschaus, E. (1986). Maternal-effect mutations altering the anterior- posterior pattern of the Drosophila embryo. Roux’s Arch. Dev. Biol. 195, 302–317. 10.1007/bf00376063.

58. Jongens, T.A., Hay, B.A., Jan, L.Y., and Jan, Y.N. (1992). The germ cell-less gene product: A posteriorly localized component necessary for germ cell development in Drosophila. Cell 70, 569–584. 10.1016/0092-8674(92)90427-e.

59. Jongens, T.A., Ackerman, L.D., Swedlow, J.R., Jan, L.Y., and Jan, Y.N. (1994). Germ cell- less encodes a cell type-specific nuclear pore-associated protein and functions early in the germ-cell specification pathway of Drosophila. Genes Dev. 8, 2123–2136. 10.1101/gad.8.18.2123.

60. Blondel, L., Besse, S., Rivard, E.L., Ylla, G., and Extavour, C.G. (2021). Evolution of a cytoplasmic determinant: Evidence for the biochemical basis of functional evolution of the novel germ line regulator Oskar. Mol. Biol. Evol. 38, 5491–5513. 10.1093/molbev/msab284.

61. Liang, L., Diehl-Jones, W., and Lasko, P. (1994). Localization of vasa protein to the Drosophila pole plasm is independent of its RNA-binding and helicase activities. Development 120, 1201–1211.

62. Johnston, D.S., Beuchle, D., and Nüsslein-Volhard, C. (1991). staufen, a gene required to localize maternal RNAs in the Drosophila egg. Cell 66, 51–63. 10.1016/0092-8674(91)90138-o.

63. Lasko, P.F., and Ashburner, M. (1990). Posterior localization of vasa protein correlates with, but is not sufficient for, pole cell development. Gene Dev 4, 905–921. 10.1101/gad.4.6.905.

64. Lehmann, R. (1992). Germ-plasm formation and germ-cell determination in Drosophila. Curr. Opin. Genet. Dev. 2, 543–549. 10.1016/s0959-437x(05)80169-8.

65. Kistler, K.E., Trcek, T., Hurd, T.R., Chen, R., Liang, F.-X., Sall, J., Kato, M., and Lehmann, R. (2018). Phase transitioned nuclear Oskar promotes cell division of Drosophila primordial germ cells. eLife 7, e37949. 10.7554/elife.37949.

66. Curnutte, H.A., Lan, X., Sargen, M., Ieong, S.M.A., Campbell, D., Kim, H., Liao, Y., Lazar, S.B., and Trcek, T. (2023). Proteins rather than mRNAs regulate nucleation and persistence of Oskar germ granules in Drosophila. Cell Rep. 42, 112723. 10.1016/j.celrep.2023.112723.

67. García-Castellanos, R., Mallorquí-Fernández, G., Marrero, A., Potempa, J., Coll, M., and Gomis-Rüth, F.X. (2004). On the Transcriptional Regulation of Methicillin Resistance. J. Biol. Chem. 279, 17888–17896. 10.1074/jbc.m313123200.

68. Jumper, J., Evans, R., Pritzel, A., Green, T., Figurnov, M., Ronneberger, O., Tunyasuvunakool, K., Bates, R., Žídek, A., Potapenko, A., et al. (2021). Highly accurate protein structure prediction with AlphaFold. Nature 596, 583–589. 10.1038/s41586-021-03819-2.

69. Mirdita, M., Schütze, K., Moriwaki, Y., Heo, L., Ovchinnikov, S., and Steinegger, M. (2022). ColabFold: making protein folding accessible to all. Nat. Methods 19, 679–682. 10.1038/s41592-022-01488-1.

70. Katoh, K., and Standley, D.M. (2013). MAFFT Multiple Sequence Alignment Software Version 7: Improvements in Performance and Usability. Mol. Biol. Evol. 30, 772–780. 10.1093/molbev/mst010.

71. Quan, J., and Tian, J. (2009). Circular Polymerase Extension Cloning of Complex Gene Libraries and Pathways. Plos One 4, e6441. 10.1371/journal.pone.0006441.

72. Quan, J., and Tian, J. (2011). Circular polymerase extension cloning for high-throughput cloning of complex and combinatorial DNA libraries. Nat Protoc 6, 242–251. 10.1038/nprot.2010.181.

73. Tanaka, T., and Nakamura, A. (2008). The endocytic pathway acts downstream of Oskar in,Drosophila,germ plasm assembly. Development 135, 1107–1117. 10.1242/dev.017293.

74. Ni, J.-Q., Liu, L.-P., Binari, R., Hardy, R., Shim, H.-S., Cavallaro, A., Booker, M., Pfeiffer, B.D., Markstein, M., Wang, H., et al. (2009). A Drosophila Resource of Transgenic RNAi Lines for Neurogenetics. Genetics 182, 1089–1100. 10.1534/genetics.109.103630.

75. Ni, J.-Q., Zhou, R., Czech, B., Liu, L.-P., Holderbaum, L., Yang-Zhou, D., Shim, H.-S., Tao, R., Handler, D., Karpowicz, P., et al. (2011). A genome-scale shRNA resource for transgenic RNAi in Drosophila. Nat. Methods 8, 405–407. 10.1038/nmeth.1592.

76. Gibson, D.G., Young, L., Chuang, R.-Y., Venter, J.C., Hutchison, C.A., and Smith, H.O. (2009). Enzymatic assembly of DNA molecules up to several hundred kilobases. Nat. Methods 6, 343–345. 10.1038/nmeth.1318.

77. DiMaio, F., Terwilliger, T.C., Read, R.J., Wlodawer, A., Oberdorfer, G., Wagner, U., Valkov, E., Alon, A., Fass, D., Axelrod, H.L., et al. (2011). Improved molecular replacement by density- and energy-guided protein structure optimization. Nature 473, 540–543. 10.1038/nature09964.

78. Adams, P.D., Grosse-Kunstleve, R.W., Hung, L., Ioerger, T.R., McCoy, A.J., Moriarty, N.W., Read, R.J., Sacchettini, J.C., Sauter, N.K., and Terwilliger, T.C. (2002). PHENIX: building new software for automated crystallographic structure determination. Acta Crystallogr. Sect. D 58, 1948–1954. 10.1107/s0907444902016657.

79. Adams, P.D., Gopal, K., Grosse-Kunstleve, R.W., Hung, L., Ioerger, T.R., McCoy, A.J., Moriarty, N.W., Pai, R.K., Read, R.J., Romo, T.D., et al. (2004). Recent developments in the PHENIX software for automated crystallographic structure determination. J. Synchrotron Radiat. 11, 53–55. 10.1107/s0909049503024130.

80. Webb, B., and Sali, A. (2016). Comparative Protein Structure Modeling Using MODELLER. Curr. Protoc. Bioinform. 54, 5.6.1–5.6.37. 10.1002/cpbi.3.

81. Söding, J., Biegert, A., and Lupas, A.N. (2005). The HHpred interactive server for protein homology detection and structure prediction. Nucleic Acids Res 33, W244–W248. 10.1093/nar/gki408.

82. McCoy, A.J., Grosse-Kunstleve, R.W., Adams, P.D., Winn, M.D., Storoni, L.C., and Read, R.J. (2007). Phaser crystallographic software. J. Appl. Crystallogr. 40, 658–674. 10.1107/s0021889807021206.

83. Emsley, P., and Cowtan, K. (2004). Coot: model-building tools for molecular graphics. Acta Crystallogr. Sect. D: Biol. Crystallogr. 60, 2126–2132. 10.1107/s0907444904019158.

84 Schrödinger, and LLC The PyMOL Molecular Graphics System, Version 3.0.

