## Supplemental Information for "The LOTUS domain of Oskar promotes localisation of both protein and mRNA components of *Drosophila* germ plasm"

This Supplemental Information contains the following elements:

- Supplementary Figure S1 through S4
- Supplementary Tables S1 through S5
- Key Reagents Table

### SUPPLEMENTAL FIGURES AND LEGENDS

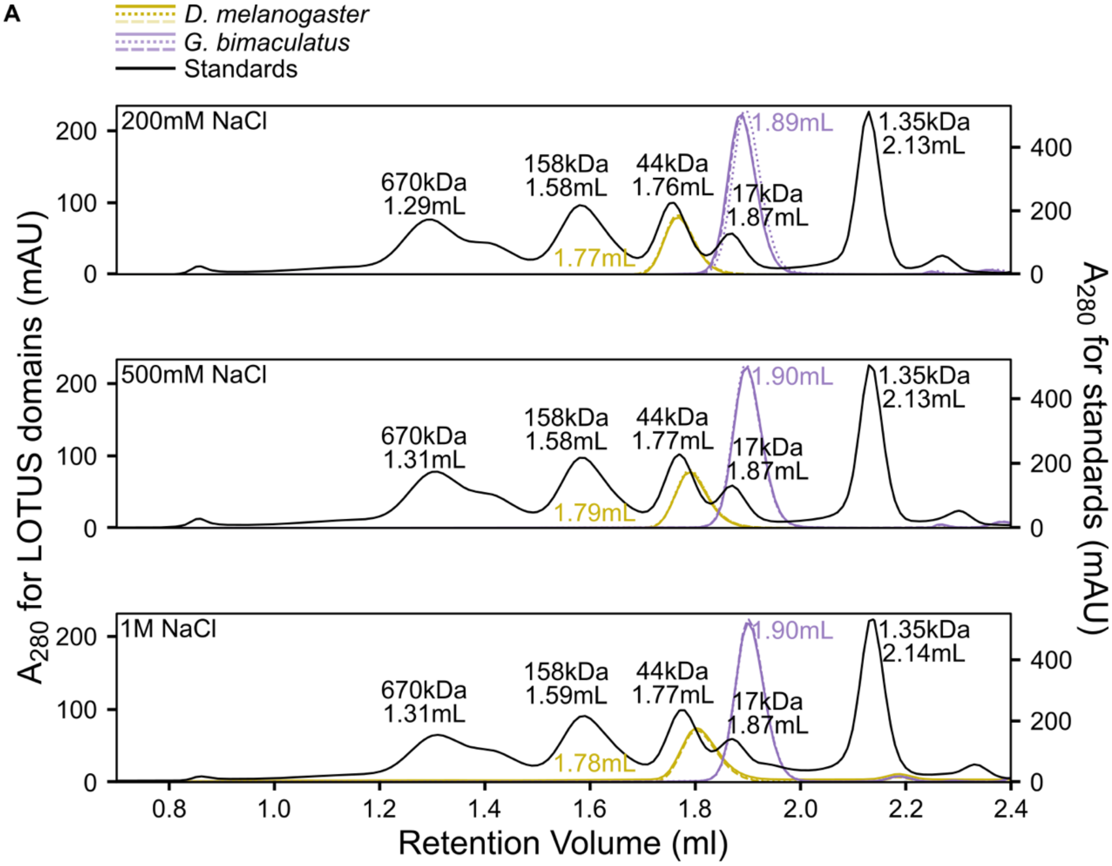


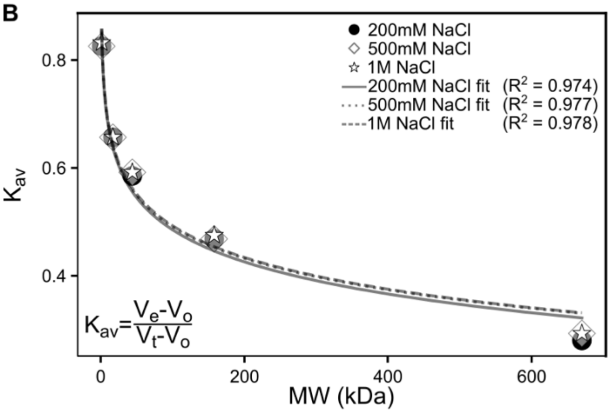


C

| [NaCl] (Mm) | *D. melanogaster* apparent MW (kDa) FW 12.3kDa | *G. bimaculatus* apparent MW (kDa) FW 10.5kDa |
|---|---|---|
| 200 | 28.8 | 11.7 |
| 500 | 26.0 | 11.2 |
| 1000 | 25.6 | 11.4 |

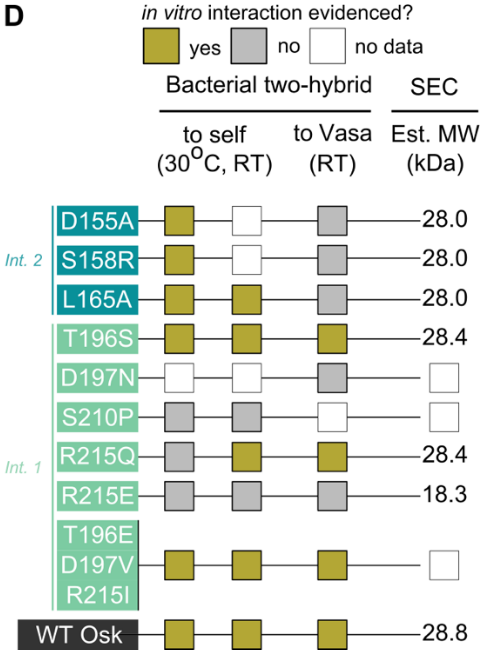

**Figure S1. *D. melanogaster* Oskar LOTUS behaves as a dimer and *G. bimaculatus* Oskar LOTUS as a monomer *in vitro*.** (B) Overlaid A280 chromatograms from triplicate gel filtration experiments of *D. melanogaster* Oskar LOTUS domain (gold curves; formula weight 12.3 kDa) and *G. bimaculatus* Oskar LOTUS domain (purple curves; formula weight 10.5 kDa) with standards (black curve; corresponding to bovine thyroglobulin (670 kDa), bovine gamma-globulin (158 kDa), chicken ovalbumin (44 kDa), horse myoglobin (17 kDa), and vitamin B12 (1.35 kDa)) performed with 20mM Tris pH 8.0 and either 200mM, 500mM, or 1M NaCl running buffer. Solid, dotted, and dashed lines represent independent runs. The monomeric molecular weights and peak retention volumes of standards and samples are indicated. All runs were performed on a Superose 6 PC 3.2/30 column (GE Healthcare). (**B**) Calibration curves for each analytical SEC buffer condition, generated by fitting the gel filtration standards data in (A). See Materials & Methods for full description of the quantification. (**C**) The apparent molecular weights of the *D. melanogaster* and *G. bimaculatus* Oskar LOTUS domains were determined by comparing their observed partition coefficients (see Materials & Methods) with the corresponding standards calibration curve in (B). The formula weight (FW) for each monomeric protein is indicated. **(D)** Bacterial two-hybrid assay (BACTH) at $30^{\circ}$C and room temperature (RT) evidencing *in vitro* interaction (gold square), lack of *in vitro* interaction (grey squares), or no data (white squares). D197N, S210P, and T196E-D197V-R215I were insoluble. Analytical SEC used to calculate estimated molecular weight (Est. MW).

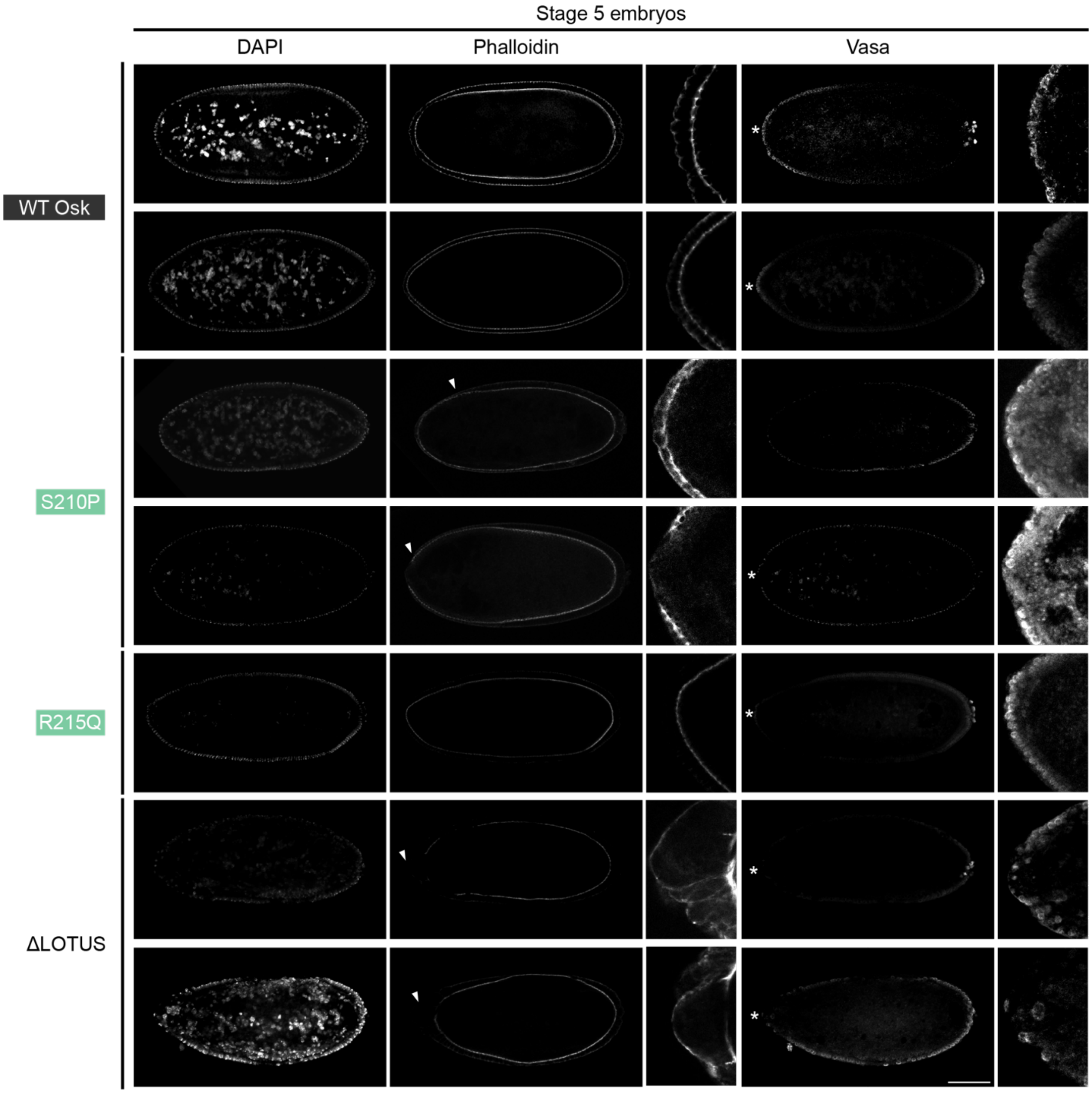


**Figure S2. Examples of anterior pole cell attempts.** Optical sections and magnified insets of confocal micrographs of stage 5 embryos from *$y^1$ $v^1$/$w^*$; P{$y^+$ $v^+$ = (oskar variant-HA-bcd3'UTR)}attp40/+; P{$w^{+mC}$=matalpha4-GAL-VP16}V37/+* mothers. Fixed embryos were stained for nuclei (DAPI), F-actin (phalloidin) for staging, the HA epitope (anti-HA antibody), and Vasa (anti-Vasa antibody) to mark pole cells. Anterior is to the left. The R215Q and last ΔLOTUS embryos were masked for visualization purposes. Micrographs were selected to show examples of phenotypes scored as developmental defects (arrowheads, shown at higher magnification to the right) and/or anterior pole cell attempts (asterisks, shown at higher magnification to the right). See Table S2 for full scoring table. Scale bar = 100μm and applies to all panels.

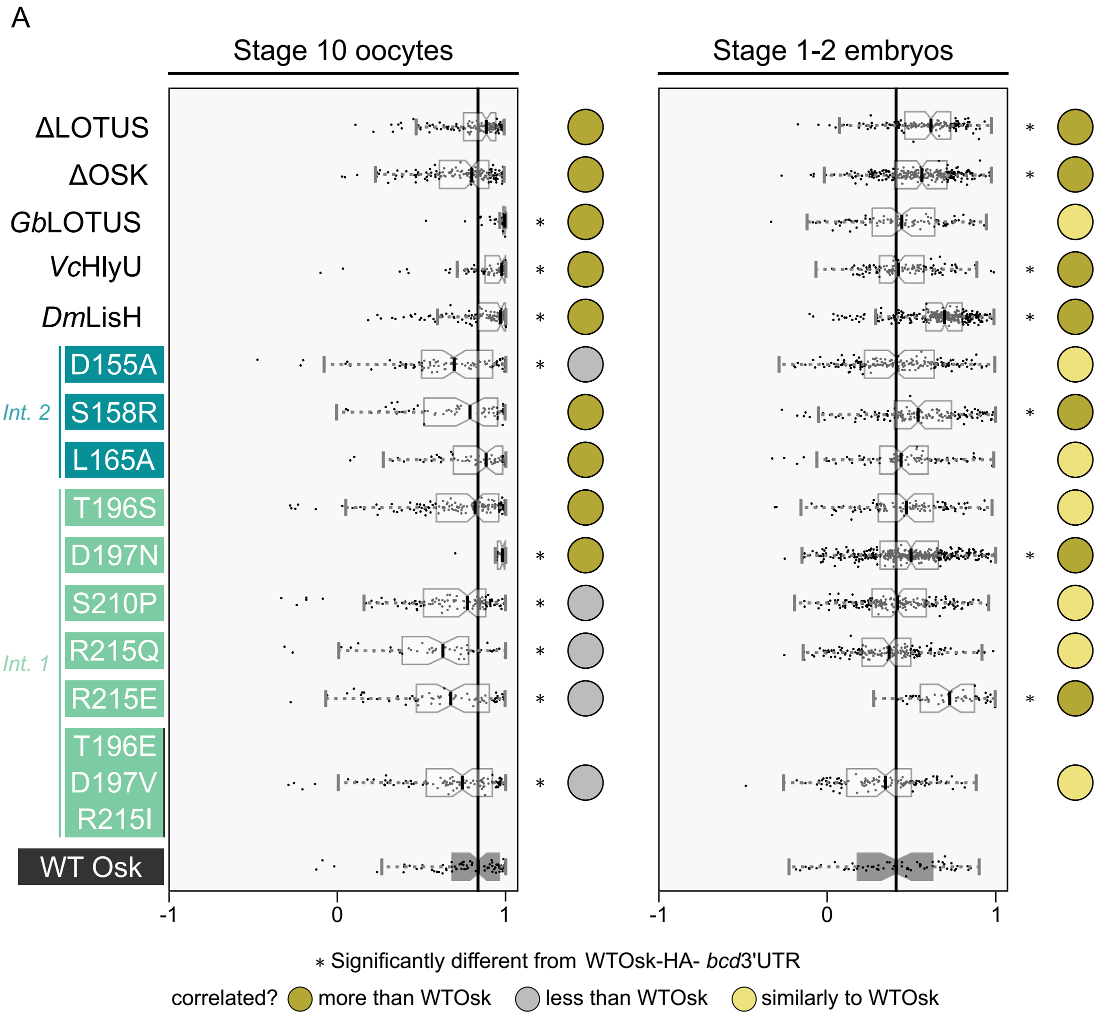
A
Stage 10 oocytes
Stage 1-2 embryos
ΔLOTUS
ΔOSK
GbLOTUS
VcHlyU
DmLisH
D155A
S158R
L165A
Int. 2
T196S
D197N
S210P
R215Q
R215E
T196E
D197V
R215I
Int. 1
WT Osk
-1
0
1
∗ Significantly different from WTOsk-HA- bcd3'UTR
correlated?
more than WTOsk
less than WTOsk
similarly to WTOsk

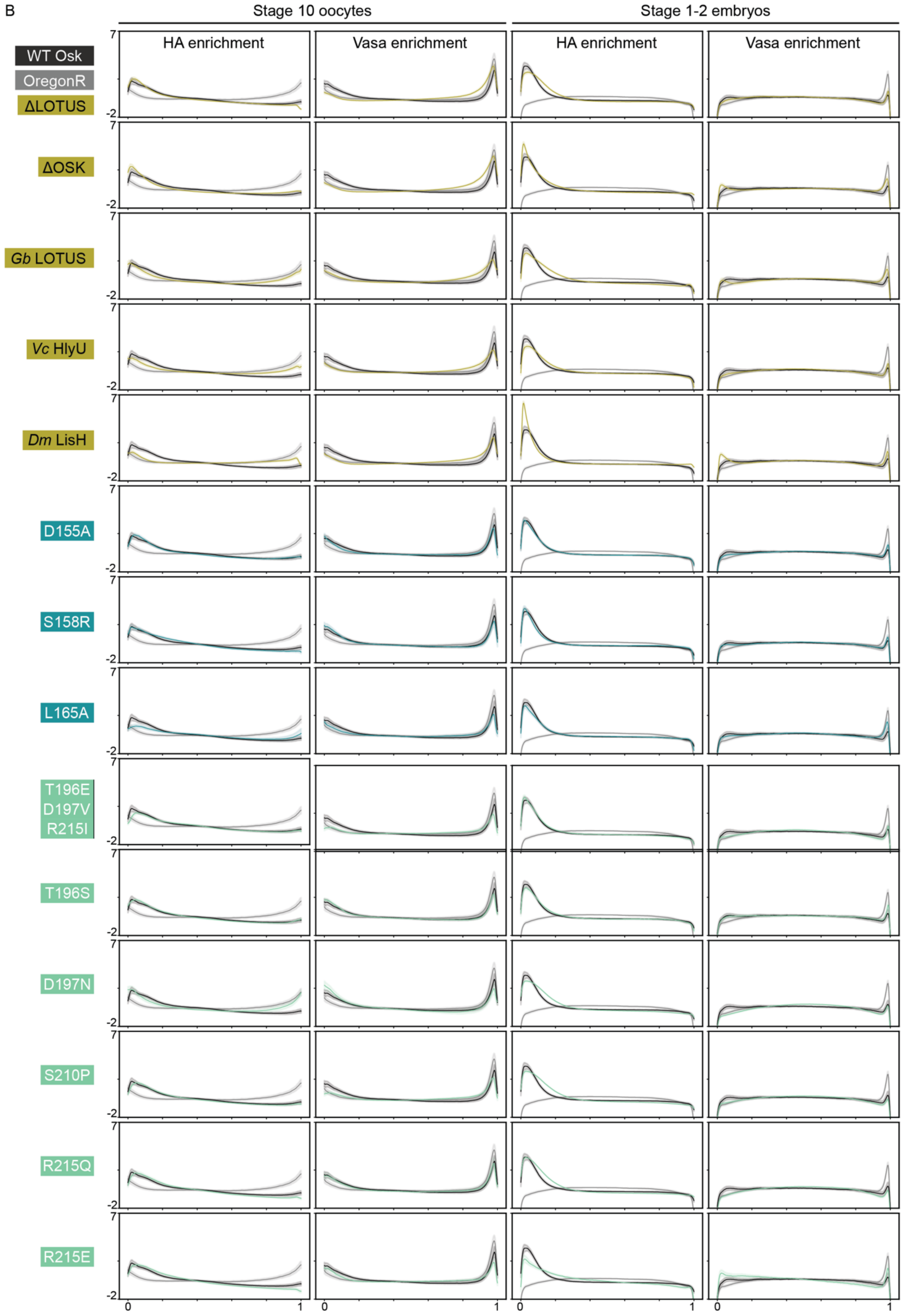

B
Stage 10 oocytes
Stage 1-2 embryos
HA enrichment
Vasa enrichment
HA enrichment
Vasa enrichment
WT Osk
OregonR
ΔLOTUS
ΔOSK
Gb LOTUS
Vc HlyU
Dm LisH
D155A
S158R
L165A
T196E
D197V
R215I
T196S
D197N
S210P
R215Q
R215E
7
-2
0
1


51

**Figure S3. Pearson correlation of HA-Vasa signal intensity suggests Oskar-Vasa anterior colocalisation in LOTUS variants. (A)** Box plots of the Pearson correlation between the HA and Vasa signal in the anterior pixels with the highest HA intensity values (with intensity values above the threshold of mean + 2 standard deviations of HA intensity values for each micrograph). Stage 10 oocytes (left) and stage 1-2 embryos (right), *y*$^{1}$ *v*$^{1}$*/w*$^{*}$*; P{y*$^{+}$ *v*$^{+}$ *= (wt-osk-HA-bcd3'UTR)}attp40/+; P{w*$^{+mC}$*=matalpha4-GAL-VP16}V37/+* (positive control) in black, median as solid black line. Significant difference from positive control distribution (p-value < 0.05, estimated using a bootstrapped test (10,000 iterations)) indicated by asterisk. Circles indicate data interpretation: HA-Vasa correlation in variant significantly more than (dark gold), less than (grey), or not significantly different from (light gold) positive control. (**B**) HA and Vasa signal enrichment (z-score) across normalised anteroposterior axis (anterior at 0, posterior at 1) in stage 10 oocytes (left) and stage 1-2 embryos (right). Mean (solid line) and 95% confidence intervals (shaded regions) shown, Oregon R (negative control) in gray, *y*$^{1}$ *v*$^{1}$*/w*$^{*}$*; P{y*$^{+}$ *v*$^{+}$ *= (wt-osk-HA-bcd3'UTR)}attp40/+; P{w*$^{+mC}$*=matalpha4-GAL-VP16}V37/+* (positive control) in black.

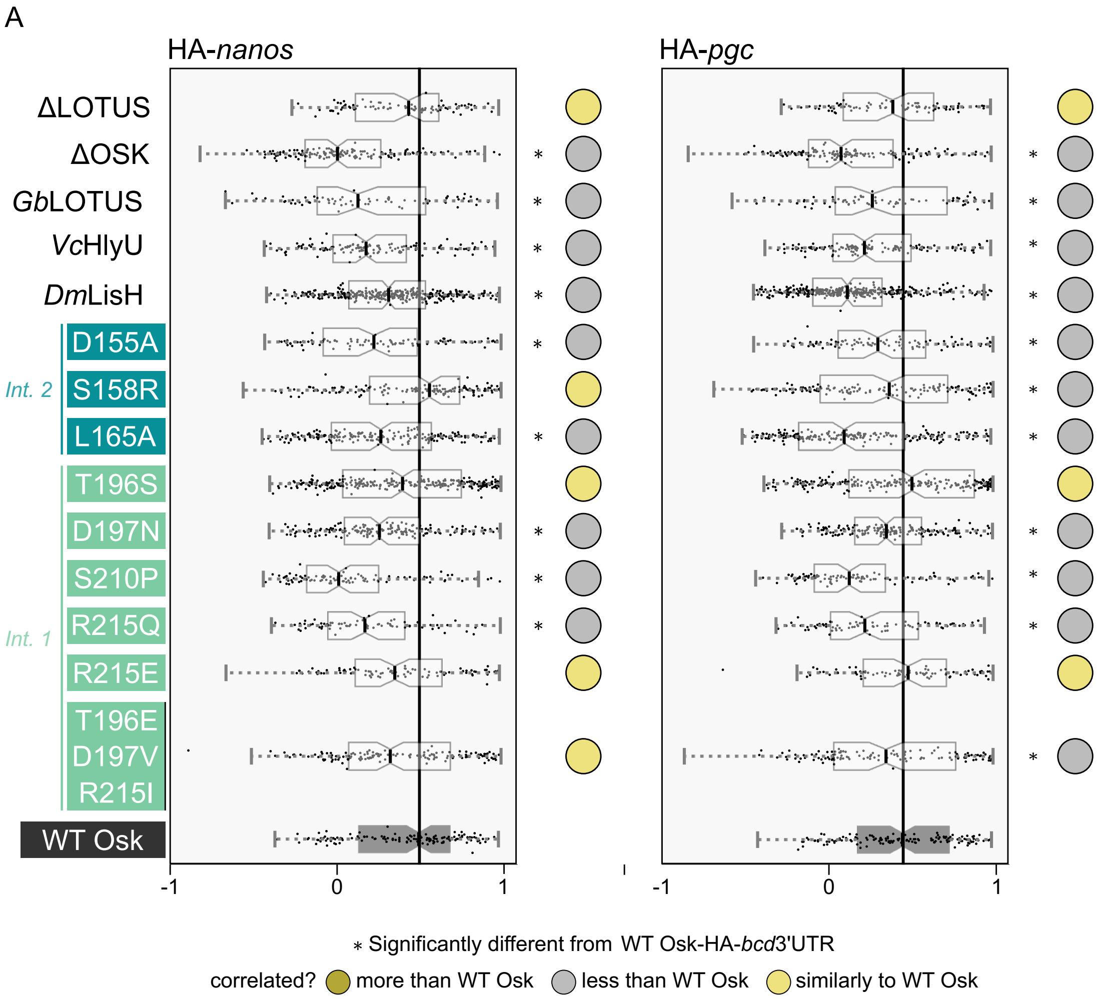

A
HA-nanos
HA-pgc
ΔLOTUS
ΔOSK
GbLOTUS
VcHlyU
DmLisH
Int. 2
D155A
S158R
L165A
Int. 1
T196S
D197N
S210P
R215Q
R215E
T196E
D197V
R215I
WT Osk
-1
0
1
* Significantly different from WT Osk-HA-bcd3'UTR
correlated? more than WT Osk less than WT Osk similarly to WT Osk

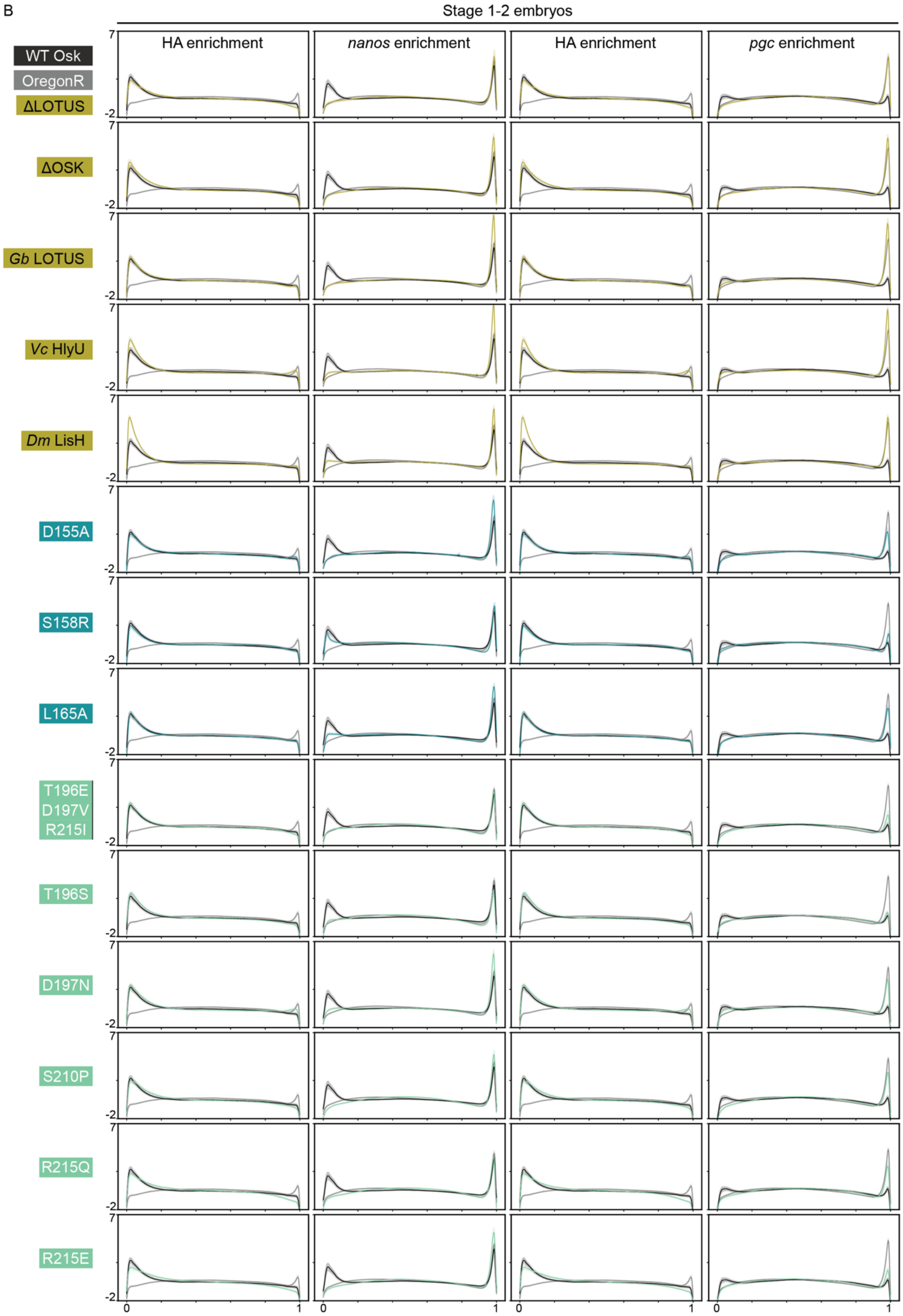

B
Stage 1-2 embryos
HA enrichment
*nanos* enrichment
HA enrichment
*pgc* enrichment
WT Osk
OregonR
ΔLOTUS
ΔOSK
*Gb* LOTUS
*Vc* HlyU
*Dm* LisH
D155A
S158R
L165A
T196E
D197V
R215I
T196S
D197N
S210P
R215Q
R215E
7
-2
0
1

**Figure S4. Pearson correlation of HA-*nanos* and HA-*pgc* signal intensity suggests Oskar-mRNA anterior colocalisation in LOTUS variants. (A)** Box plots of the Pearson correlation between the HA and *nanos* (left) and *pgc* (right) signals in the anterior pixels with the highest HA intensity values (with intensity values above the threshold of mean + 2 standard deviations of HA intensity values for each micrograph). *y*$^{1}$ *v*$^{1}$*/w*$^{*}$*; P{y*$^{+}$ *v*$^{+}$ *= (wt-osk-HA-bcd3'UTR)}attp40/+; P{w*$^{+mC}$*=matalpha4-GAL-VP16}V37/+* (positive control) in black, median as solid black line ar. Significant difference from positive control distribution (p-value < 0.05, estimated using a bootstrapped test (10,000 iterations)) indicated by asterisk. Circles indicate interpretation of data: HA-mRNA correlation in variant significantly more than (dark gold), less than (grey), or not significantly different from (light gold) *y*$^{1}$ *v*$^{1}$*/w*$^{*}$*; P{y*$^{+}$ *v*$^{+}$ *= (wt-osk-HA-bcd3'UTR)}attp40/+; P{w*$^{+mC}$*=matalpha4-GAL-VP16}V37/+*. (**B**) HA and *nanos* (left) and HA and *pgc* (right) signal enrichment (z-score) across normalized anteroposterior axis (anterior at 0, posterior at 1) in stage 1-2 embryos. Mean (solid line) and 95% confidence intervals (shaded regions) shown, Oregon R (negative control) in gray, *y*$^{1}$ *v*$^{1}$*/w*$^{*}$*; P{y*$^{+}$ *v*$^{+}$ *= (wt-osk-HA-bcd3'UTR)}attp40/+; P{w*$^{+mC}$*=matalpha4-GAL-VP16}V37/+* (positive control) in black.

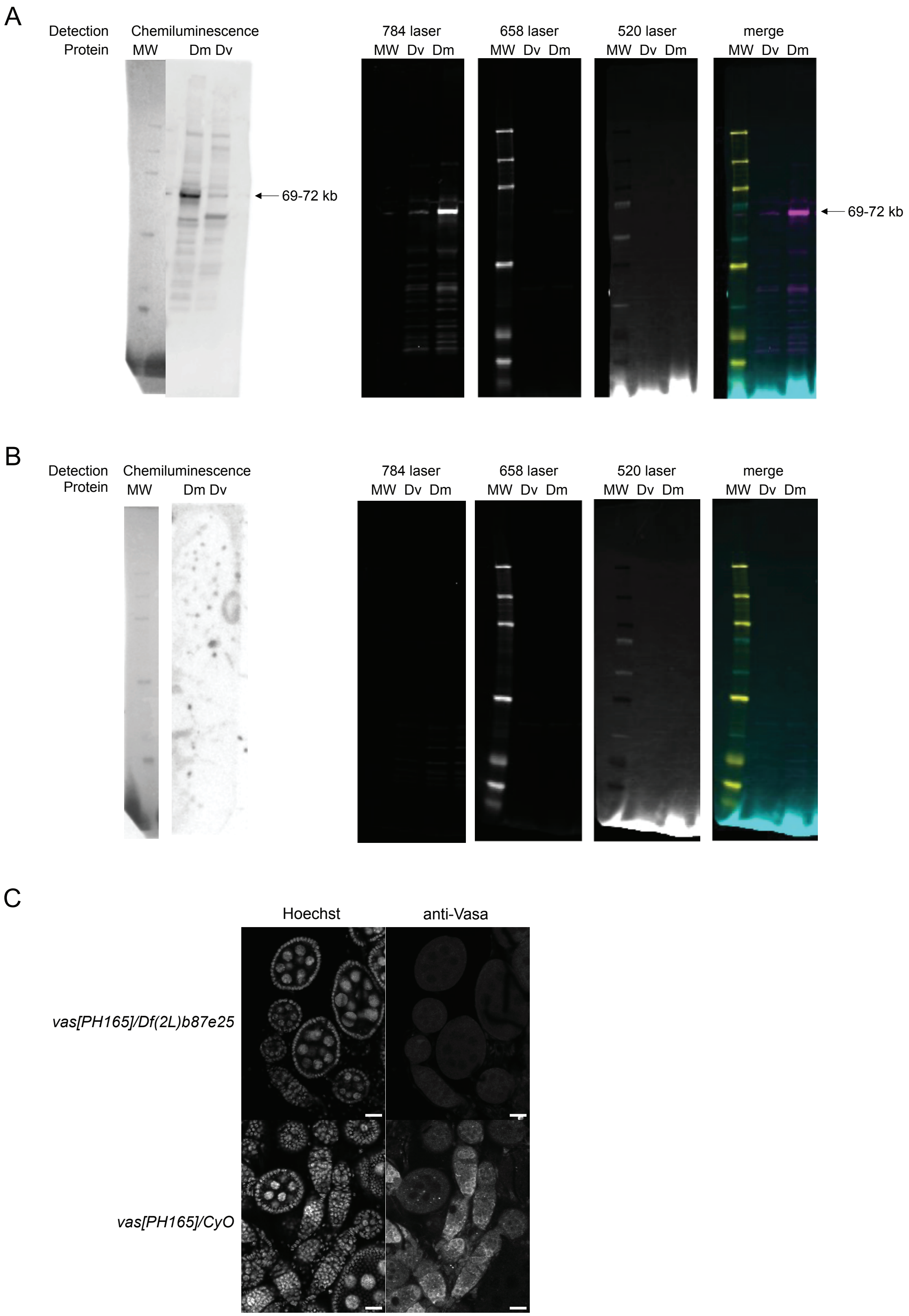
A
Detection
Protein
Chemiluminescence
MW
Dm Dv
69-72 kb
784 laser
MW Dv Dm
658 laser
MW Dv Dm
520 laser
MW Dv Dm
merge
MW Dv Dm
69-72 kb
B
Detection
Protein
Chemiluminescence
MW
Dm Dv
784 laser
MW Dv Dm
658 laser
MW Dv Dm
520 laser
MW Dv Dm
merge
MW Dv Dm
C
Hoechst
anti-Vasa
vas[PH165]/Df(2L)b87e25
vas[PH165]/CyO

**Figure S5.** *D. melanogaster* anti-Vasa antibody validation. (**A–B)** Chemiluminescent (left) and fluorescent (right) Western Blots with *D. melanogaster* (Dm) and *D. virilis* (Dv) ovary lysates. Lanes with molecular weights are marked MW. Chemiluminescent Western blots were imaged using a CCD camera on an Azure Sapphire Biomolecular Imager. Fluorescent blots were excited by three laser lines and captured using APD detectors on an Azure Sapphire Biomolecular Imager. Signal from the primary antibody is excited by the 784 nm laser, while the molecular weight is excited by the 658 nm and 520 nm lasers. Bands at expected product sizes are marked with an arrow (*D. melanogaster* Vasa ~72 kb, *D. virilis* Vasa ~69kb). Blots were incubated in anti-*D. melanogaster* Vasa (A) or no primary antibody (B). **(C)** Vasa antibody staining in *D. melanogaster* ovaries in flies with genotypes *vas[PH165]/Df(2L)b87e25* (*vasa* null) and *vas[PH165]/CyO* (control). Images are single optical sections and scale bar represents 20 µm.

**SUPPLEMENTAL TABLES AND LEGENDS**

**Table S1. Summary of diffraction data for Losk139-241 crystal forms 1–4**

| Crystal Form | 1 | 2 | 3 | 4 |
|---|---|---|---|---|
| **Sample** | Losk139-241 | Losk139-241 | Losk139-241 | Losk139-241 + $HgCl_2$ |
| **Space Group** | P4 | I4 | C222 | $P12_11$ |
| **a, b, c (Å)** | 138.95, 138.95, 61.90 | 141.1, 141.1, 219.9 | 128.6, 205.4, 172.4 | 39.60, 63.57, 40.15 |
| **α, β, γ (º)** | 90, 90, 90 | 90, 90, 90 | 90, 90, 90 | 90, 97.22, 90 |
| **Wavelength (Å)** | 0.9800 | 0.9800 | 0.9800 | 1.0039 |
| **No. of unique reflections** | 18398 | 20643 | 18656 | 13341 |
| **Resolution Range (Å)** | 50 – 3.21 | 50 – 3.13 | 50 – 4.16 | 50 – 2.00 |
| **$R_{merge}$** | 0.053 | 0.107 | 0.158 | 0.198 |
| **Overall $I/\sigma(I)$** | 8.0 (1.2) | 9.6 (1.3) | 5.4 (1.1) | 16.0 (1.9) |
| **Completeness (%)** | 92.6 | 99.0 | 98.4 | 100.0 (100.0) |
| **Multiplicity** | 1,8 | 3.5 | 2.8 | 16.3 |
| **Anomalous completeness (%)** | - | - | - | 99.9 (99.9) |
| **Anomalous multiplicity** | - | - | - | 8.2 |

**Table S2. Summary of stage 5 embryo analysis for presence of anterior pole cells.**

| | | | | HA-positive[a] | | | HA-negative[b] | | |
|---|---|---|---|---|---|---|---|---|---|
| **Variant** | **Embryos[c]** | **Stage 5[d]** | **HA positive[e]** | **Developmental defect[f]** | **Anterior pole cell attempt[g]** | **Anterior pole cells[h]** | **Developmental defect[f]** | **Anterior pole cell attempt[g]** | **Anterior pole cells[h]** |
| **Oregon R** | 347 | 298 | 4 | 2 | 0 | 0 | 1 | 0 | 0 |
| **ΔLOTUS** | 549 | 524 | 314 | 4 | 2 | 3 | 14 | 5 | 1 |
| **ΔOSK** | 222 | 217 | 216 | 23 | 24 | 0 | 1 | 1 | 0 |
| **Gb LOTUS** | 424 | 412 | 148 | 6 | 6 | 0 | 35 | 28 | 0 |
| **Vc HlyU** | 703 | 696 | 172 | 3 | 0 | 0 | 12 | 21 | 0 |
| **Dm LisH** | 443 | 436 | 372 | 36 | 11 | 0 | 16 | 8 | 0 |
| **D155A** | 920 | 874 | 307 | 28 | 0 | 0 | 16 | 1 | 2 |
| **S158R** | 501 | 469 | 363 | 8 | 1 | 6 | 2 | 0 | 0 |
| **L165A** | 742 | 689 | 379 | 44 | 0 | 2 | 33 | 0 | 0 |
| **T196S** | 601 | 586 | 384 | 42 | 4 | 8 | 10 | 0 | 1 |
| **D197N** | 586 | 549 | 307 | 24 | 2 | 1 | 34 | 0 | 0 |
| **S210P** | 853 | 799 | 355 | 48 | 2 | 2 | 32 | 2 | 8 |
| **R215Q** | 885 | 836 | 439 | 19 | 10 | 0 | 9 | 2 | 0 |
| **R215E** | 1482 | 1376 | 118 | 17 | 0 | 0 | 92 | 0 | 3 |
| **Triple*** | 427 | 417 | 200 | 2 | 2 | 0 | 3 | 1 | 0 |
| **WT Osk** | 648 | 625 | 238 | 14 | 26 | 36 | 9 | 60 | 21 |

[a] Analysis of stage 5 embryos with anterior HA enrichment.
[b] Analysis of stage 5 embryos without anterior HA enrichment, indicating either absence of HA-tagged Oskar variant at the anterior or inefficient HA staining.
[c] Number of embryos imaged per variant. See Materials & Methods for collection, staining, and imaging protocol.
[d] Number of imaged embryos visually determined to be stage 5 based on phalloidin stain showing completed cellularisation but not gastrulation.
[e] Number of stage 5 embryos visually determined to show anterior enrichment of HA signal, indicating anterior enrichment of HA-tagged Oskar variant.
[f] Number of stage 5 embryos visually determined to show developmental defects. Developmental defect phenotypes include uneven cellularisation around the periphery of the embryo and cellular budding reminiscent of pole cell budding at ectopic (non-posterior or anterior) locations. See Fig. SX for examples.
[g] Number of stage 5 embryos visually determined to show anterior PC attempts based on nuclear (DAPI) and Vasa stain. PC attempts defined as budding, high Vasa signal, or spherical nuclei in cells at the anterior, but with no fully budded PCs clearly identifiable. See Fig. 4A and Fig. S3 for examples.
[h] Number of stage 5 embryos visually determined to show anterior PCs based on nuclear (DAPI) and Vasa stain. For variants with a nonzero number of embryos with anterior pole cells, posterior and anterior pole cells counted for embryos with anterior pole cells (Fig. 4C).
[*] Triple refers to the triple mutant T196E-D197V-R215I.

**Table S3. Statistical Analysis of anterior pole cell  incidence and counts.**

| Variant | N[a] | Anterior pole cells[b] | p-value[c] (anterior pole cell incidence vs. Negative Control) | p-value[c] (anterior pole cell incidence vs. Positive Control) | p-value[d] (posterior pole cell count vs. Negative Control) | p-value[d] (anterior ole cell count vs. Positive Control) |
|---|---|---|---|---|---|---|
| **Oregon R** | 298 | 0 | | 4.144835e-14 | | 0 |
| **ΔLOTUS** | 314 | 3 | 0.134433 | 3.000754e-11 | 0.10813455 | 0.47913206 |
| **ΔOSK** | 216 | 0 | 1 | 2.043245e-11 | | |
| **Gb LOTUS** | 148 | 0 | 1 | 9.115716e-09 | | |
| **Vc HlyU** | 172 | 0 | 1 | 9.388045e-10 | | |
| **Dm LisH** | 372 | 0 | 1 | 3.395834e-16 | | |
| **D155A** | 307 | 0 | 1 | 2.230220e-14 | | |
| **S158R** | 363 | 6 | 0.026920 | 2.353295e-10 | 2.91486379e-09 | 0.00855579 |
| **L165A** | 379 | 2 | 0.313037 | 6.843532e-14 | 0 | 0.38321883 |
| **T196S** | 384 | 8 | 0.009781 | 1.093258e-09 | 1.89603991e-05 | 0.09808461 |
| **D197N** | 307 | 1 | 0.507438 | 5.065534e-13 | 1.11022302e-16 | 0.00531489 |
| **S210P** | 355 | 2 | 0.295169 | 2.850252e-13 | 0.00326256 | 5.44009282e-15 |
| **R215Q** | 439 | 0 | 1 | 7.170191e-18 | | |
| **R215E** | 118 | 0 | 1 | 1.943480e-07 | | |
| **Triple*** | 200 | 0 | 1 | 7.843703e-11 | | |
| **WT Osk** | 238 | 36 | 4.144835e-14 | | 2.25375274e-14 | |

* Triple refers to the triple mutant T196E-D197V-R215I.
[a] Number of stage 5 embryos (Oregon R) or number of stage 5 embryos with anterior HA enrichment.
[b] Number of embryos visually determined to show anterior PCs.
[c] p-value (calculated using Fisher's exact test, based on a 2×2 contingency table; exact probabilities are computed without relying on large-sample approximations).
[d] p-value (estimated using a bootstrapped test (10,000 iterations), comparing the observed difference in means from control distribution to a distribution generated by resampling with replacement.

| Table S4. Statistical Analysis of Vasa enrichment in oocytes and embryos. | | | | | | | | |
|---|---|---|---|---|---|---|---|---|
| | Stage 10 Oocytes | | | | Stage 1-2 embryos | | | |
| **Variant** | **Count[a]** | **p-value[b] (Integrated Anterior Enrichment vs. Positive Control)** | **p-value[b] (Integrated Anterior Enrichment vs. Negative Control)** | **p-value[b] (Pearson Correlation vs. Positive Control)** | **Count[a]** | **p-value[b] (Integrated Anterior Enrichment vs. Positive Control)[b]** | **p-value[b] (Integrated Anterior Enrichment vs. Negative Control)** | **p-value[b] (Pearson Correlation vs. Positive Control)** |
| **Oregon R** | 84 | 6.55018591e-07 | | | 189 | 0.00124249 | | |
| **ΔLOTUS** | 138 | 5.89546013e-17 | 0.04460279 | 0.08094359 | 160 | 0.14182322 | 0.00535144 | 1.02738993e-06 |
| **ΔOSK** | 150 | 2.03899506e-18 | 0.02280415 | 0.10769372 | 265 | 0.38275709 | 9.17341758e-11 | 5.73561552e-05 |
| **Gb LOTUS** | 78 | 1.08245031e-08 | 0.33320252 | 2.21267449e-13 | 100 | 0.05963263 | 0.0152061 | 0.12304134 |
| **Vc HlyU** | 109 | 9.95357718e-13 | 0.19365831 | 1.91191839e-05 | 122 | 0.01947077 | 0.13663505 | 0.04612047 |
| **Dm LisH** | 182 | 5.56884297e-17 | 0.05777246 | 7.75637362e-05 | 307 | 0.18007473 | 1.73749903e-13 | 1.15651932e-12 |
| **D155A** | 97 | 0.01140579 | 0.00051423 | 0.00258703 | 191 | 0.01958909 | 0.11748532 | 0.19789815 |
| **S158R** | 66 | 0.02609067 | 3.78562737e-11 | 0.07952136 | 154 | 0.13775126 | 0.00311394 | 0.00022194 |
| **L165A** | 69 | 0.01265417 | 0.0076116 | 0.20661084 | 92 | 0.20000143 | 0.00187369 | 0.1183661 |
| **T196S** | 150 | 0.11665846 | 6.63691324e-13 | 0.09257797 | 118 | 0.48573134 | 0.00014762 | 0.09177468 |
| **D197N** | 31 | 0.0148029 | 8.1539997e-11 | 3.55437901e-12 | 409 | 3.68680593e-06 | 0.00531154 | 0.006517 |
| **S210P** | 126 | 1.95206258e-06 | 0.13607942 | 0.00450611 | 210 | 0.00954832 | 0.20107368 | 0.18511517 |
| **R215Q** | 50 | 0.13839737 | 1.93235927e-07 | 4.85141466e-05 | 179 | 0.00010435 | 0.10580469 | 0.20384861 |
| **R215E** | 73 | 0.01924888 | 0.00041599 | 0.00084778 | 58 | 0.00206767 | 7.92366173e-13 | 1.78301818e-13 |
| **Triple[*]** | 105 | 0.00014175 | 0.12874218 | 0.00619258 | 127 | 0.00437658 | 0.41212536 | 0.07212177 |
| **WT Osk** | 85 | | 7.8202681e-07 | | 65 | | 0.00141613 | |

[*] Triple refers to the triple mutant T196E-D197V-R215I.
[a] Number of oocytes or embryos visually determined to show anterior enrichment of HA signal, indicating anterior enrichment of HA-tagged Oskar variant.
[b] p-value (difference from control distribution, 10,000 Simulations, bootstrapping with resampling)

| **Table S5. Statistical Analysis of *nanos* and *pgc* enrichment in stage 1-2 embryos.** | | | | | | | | |
|---|---|---|---|---|---|---|---|---|
| | ***nanos*** | | | | ***pgc*** | | | |
| **Variant** | **Count[a]** | **p-value[b] (Integrated Anterior Enrichment vs. Positive Control)** | **p-value[b] (Integrated Anterior Enrichment vs. Negative Control)** | **p-value[b] (Pearson Correlation vs. Positive Control)** | **Count[a]** | **p-value[b] (Integrated Anterior Enrichment vs. Positive Control)[b]** | **p-value[b] (Integrated Anterior Enrichment vs. Negative Control)** | **p-value[b] (Pearson Correlation vs. Positive Control)** |
| **Oregon R** | 723 | 1.51552666e-31 | | | 592 | 4.4135902e-08 | | |
| **ΔLOTUS** | 85 | 3.17469149e-20 | 0.03501306 | 0.28582801 | 85 | 2.72834324e-17 | 3.19958328e-10 | 0.06045782 |
| **ΔOSK** | 150 | 2.34761833e-44 | 4.77285821e-09 | 6.3090931e-20 | 150 | 1.68497567e-16 | 6.31120934e-11 | 1.51666976e-15 |
| **Gb LOTUS** | 73 | 5.10266219e-37 | 0.00027754 | 1.2854597e-05 | 73 | 0.00382874 | 0.05428115 | 0.00657417 |
| **Vc HlyU** | 130 | 1.90779859e-18 | 5.4366558e-07 | 4.89546054e-07 | 130 | 0.1235291 | 4.02455846e-13 | 3.09157442e-06 |
| **Dm LisH** | 362 | 1.54521935e-09 | 1.22124533e-15 | 0.00050271 | 362 | 0.00668283 | 5.3265015e-05 | 1.63376578e-21 |
| **D155A** | 90 | 1.51706518e-27 | 0.30760755 | 0.00014871 | 90 | 5.15788935e-05 | 0.41437526 | 0.01011794 |
| **S158R** | 158 | 0.01816407 | 0. | 0.08821007 | 156 | 0.00486035 | 0.00503855 | 0.00912196 |
| **L165A** | 215 | 1.41928405e-09 | 1.76503256e-12 | 0.00011704 | 215 | 1.02274157e-07 | 0.25075921 | 5.50798673e-15 |
| **T196S** | 289 | 1.2330628e-27 | 0.27089413 | 0.2882354 | 245 | 2.48909122e-09 | 0.04313497 | 0.26528852 |
| **D197N** | 173 | 2.73857579e-12 | 4.85779973e-08 | 0.00011259 | 170 | 0.19026173 | 6.25437357e-09 | 0.01049373 |
| **S210P** | 98 | 1.29226955e-60 | 1.24171548e-24 | 6.43372641e-13 | 98 | 2.15858897e-22 | 1.0067283e-18 | 1.04211795e-09 |
| **R215Q** | 79 | 2.43610099e-42 | 1.68498198e-09 | 1.06647914e-06 | 79 | 6.69470709e-12 | 3.97562825e-05 | 0.00013527 |
| **R215E** | 88 | 2.46350993e-49 | 1.5330683e-13 | 0.17277567 | 85 | 1.92576858e-06 | 0.12558829 | 0.36006112 |
| **Triple[*]** | 144 | 7.71456903e-30 | 0.03886315 | 0.11417878 | 125 | 7.20415414e-18 | 1.43887471e-11 | 0.019827 |
| **WT Osk** | 142 | | 0. | | 137 | | 5.62784721e-08 | |

[*] Triple refers to the triple mutant T196E-D197V-R215I.

[a] Number of oocytes or embryos visually determined to show anterior enrichment of HA signal, indicating anterior enrichment of HA-tagged Oskar variant.

[b] p-value (Difference from Control Distribution, 10,000 Simulations, bootstrapping with resampling)

**KEY RESOURCES TABLE**

| REAGENT or RESOURCE | SOURCE | IDENTIFIER |
|---|---|---|
| **Antibodies** | | |
| Mouse anti HA-Tag monoclonal antibody | ABclonal | AE008 |
| Rat anti-HA High Affinity monoclonal antibody | Roche | 3F10 |
| Rabbit anti Vasa polyclonal antibody | Laboratory of Prashanth Rangan (Icahn School of Medicine at Mount Sinai) | |
| Chicken anti Vasa polyclonal antibody | This paper | 9276 |
| DAPI, for nucleic acid staining | Sigma-Aldrich | D9542 |
| Rhodamine Phalloidin | Thermo Fisher Scientific | R415 |
| Goat anti-Mouse IgG (H+L) Cross-Adsorbed Secondary Antibody, Alexa Fluor™ 488 | Thermo Fisher Scientific | A-11001 |
| Goat anti-Rat IgG (H+L) Cross-Adsorbed Secondary Antibody, Alexa Fluor™ 488 | Thermo Fisher Scientific | A-11006 |
| Goat anti-Rabbit IgG (H+L) Highly Cross-Adsorbed Secondary Antibody, Alexa Fluor™ 647 | Thermo Fisher Scientific | A-21245 |
| Goat anti-Chicken IgY (H+L) Cross-Adsorbed Secondary Antibody, Alexa Fluor™ 633 | Thermo Fisher Scientific | A-21103 |
| HCR™ RNA-FISH anti-nanos probe (v3.0) | Molecular Instruments | |
| HCR™ RNA-FISH anti-pgc probe (v3.0) | Molecular Instruments | |
| HCR™ RNA-FISH amplifier B5-555 (v3.0) | Molecular Instruments | |
| HCR™ RNA-FISH amplifier B4-647 (v3.0) | Molecular Instruments | |
| **Bacterial and virus strains** | | |
| BL21(DE3) *E. coli* | | |
| BTH101 *E. coli* | Euromedex | EUK001 |
| DHM1 *E. coli* | Euromedex | EUK001 |
| **Chemicals, peptides, and recombinant proteins** | | |

| | | |
|---|---|---|
| 2-mercaptoethanol, Omnipur | Avantor | Cat# EM-6050 |
| 5x Pierce Lane Marker Non-Reducing Sample Buffer | Thermo Scientific | Cat# 39001 |
| Precision Plus Protein Dual Color Standards | Bio-Rad | Cat# 1610394 |
| Precision Plus Protein WesternC Blotting Standards | Bio-Rad | Cat# 1610376 |
| SuperSignal West Pico PLUS Chemiluminescent Substrate | Thermo Scientific | Cat# 34580 |
| Sodium azide | Sigma-Aldrich | Cat# S2002 |
| Fluorescent Blot Blocking Buffer | Azure | Cat# AC2190 |
| Azure Blot Washing Buffer | Azure | Cat# AC2113 |
| *Drosophila melanogaster* Vasa (UniProt ID: M9PBB5, M1-E206 aa) | This paper | N/A |
| Ni-NTA Agarose | QIAGEN or Thermo Scientific | |
| Gel Filtration Standards | BIO-RAD | |
| Amicon Ultracel Devices | Millipore | |
| **Critical commercial assays** | | |
| Pierce Detergent Compatible Bradford Assay Kit | Thermo Scientific | Cat# 23246 |
| Trans-Blot Turbo RTA Mini 0.2 µm PVDF Transfer Kit | Bio-Rad | Cat# 1704272 |
| Bacterial Adenylate Cyclase-based Two-Hybrid (BACTH) kit | Euromedex | EUK001 |
| **Deposited data used in analyses** | | |
| Raw and analysed data | This paper | |
| Oskar protein sequence *D. melanogaster* | UniProt | Acc. No. P25158 |
| Oskar protein sequence *D. sechellia* | UniProt | Acc. No. B4HKZ1 |
| Oskar protein sequence *D. simulans* | UniProt | Acc. No. B4QXC8 |
| Oskar protein sequence *D. yakuba* | UniProt | Acc. No. |

| | | |
|---|---|---|
| | | B4PTX6 |
| Oskar protein sequence *D. immigrans* | UniProt | Acc. No. A1Y1T7 |
| Oskar protein sequence *D. virilis* | UniProt | Acc. No. Q24741 |
| Oskar protein sequence *D. pseudoobscura* | UniProt | Acc. No. Q295Q4 |
| Oskar protein sequence *D. erecta* | UniProt | Acc. No. B3P1W4 |
| Oskar protein sequence *D. ananassae* | UniProt | Acc. No. B3LZ06 |
| Oskar protein sequence *D. persimilis* | UniProt | Acc. No. B4GFV0 |
| Oskar protein sequence *D. willistoni* | UniProt | Acc. No. B4N815 |
| Oskar protein sequence *D. mojavensis* | UniProt | Acc. No. B4K9E4 |
| Oskar protein sequence *D. grimshawi* | UniProt | Acc. No. B4JTJ1 |
| Oskar protein sequence *Gryllus bimaculatus* | UniProt | Acc. No. K4MTL4 |
| *Vibrio cholera* transcription factor HlyU | UniProt | Acc. No. C3LST3 |
| *D. melanogaster lis1* LisH domain | UniProt | Acc. No. Q7KNS3 |
| Crystal structure of the LOTUS domain (aa 139-222) of *D. melanogaster* Oskar in C222 | [1] | RCSB PDB 5A49 |
| Crystal structure of the NTD L199M of *D. melanogaster* Oskar protein | [2] | RCSB PDB 5CD7 |
| Structure of the LOTUS domain of *D. melanogaster* Oskar in complex with the C-terminal RecA-like domain of *D. melanogaster* Vasa | [3] | RCSB PDB 5NT7 |
| **Experimental models: *D. melanogaster* strains** | | |
| *w[*]; P{w[+mC]=matalpha4-GAL-VP16}V37* | Bloomington Drosophila | 7063 |

| | | |
|---|---|---|
| | Stock Center | |
| *y[1] v[1]; Sco/CyO* | Laboratory of Norbert Perrimon lab (Harvard Medical School) | N/A |
| *y[1] v[1]; attP40* | Genetic Services, Inc. | |
| *y[1] v[1]; P{y[+] v[+]=(wt-osk-HA-bcd3'UTR)}attp40* | This paper | WT Osk |
| *y[1] v[1]; P{y[+]v[+]=(Δ140-241-osk-HA-bcd3'UTR)}attp40* | This paper | ΔLOTUS |
| *y[1] v[1]; P{y[+]v[+]=(Δ401-606-osk-HA-bcd3'UTR)}attp40* | This paper | ΔOSK |
| *y[1] v[1]; P{y[+]v[+]=(Gbosk2-90-Dmosk-HA-bcd3'UTR)}attp40* | This paper | Gb LOTUS |
| *y[1] v[1]; P{y[+]v[+]=(VcHlyU2-109-Dmosk-HA-bcd3'UTR)}attp40* | This paper | Vc HlyU |
| *y[1] v[1]; P{y[+]v[+]=(DmLis2-86-Dmosk-HA-bcd3'UTR)}attp40* | This paper | Dm LisH |
| *y[1] v[1]; P{y[+]v[+]=(Dmosk-D155A-HA-bcd3UTR)}attp40* | This paper | D155A |
| *y[1] v[1]; P{y[+] v[+]=(Dmosk-S158R-HA-bcd3UTR)}attp40* | This paper | S158R |
| *y[1] v[1]; P{y[+]v[+]=(Dmosk-L165A-HA-bcd3UTR)}attp40* | This paper | L165A |
| *y[1] v[1]; P{y[+] v[+]=(Dmosk-T196S-HA-bcd3UTR)}attp40* | This paper | T196S |
| *y[1] v[1]; P{y[+] v[+]=(Dmosk-D197N-HA-bcd3UTR)}attp40* | This paper | D197N |
| *y[1] v[1]; P{y[+] v[+]=(Dmosk-S210P-HA-bcd3UTR)}attp40* | This paper | S210P |
| *y[1] v[1]; P{y[+] v[+]=(Dmosk-R215Q-HA-bcd3UTR)}attp40* | This paper | R215Q |
| *y[1] v[1]; P{y[+] v[+]=(Dmosk-R215E-HA-bcd3UTR)}attp40* | This paper | R215E |
| *y[1] v[1]; P{y[+]v[+]=(Dmosk-T196E-D197V-R215I-HA-bcd3UTR)}attp40* | This paper | T196E |

| | | |
|---|---|---|
| *vas[PH165]/CyO; Drop/TM3* | Laboratory of Paul Lasko (McGill University) | N/A |
| *Df(2L)b87e25/CyO* | Bloomington Drosophila Stock Center | BDSC 3138 |
| **Recombinant DNA** | | |
| pET-43.1b(+) expression vector | Novagen | |
| pET-Losk139-241_C10H | This paper | N/A |
| pET-151-Gbosk1-440 | [4] | N/A |
| pET-151-Gbosk1-90 | This paper | N/A |
| *D. melanogaster* transgenic vector pVALIUM22 | Perrimon lab (Harvard Medical School) | N/A |
| pVAL22-Dmosk-bcd3UTR | This paper | N/A |
| pVAL22-Dmosk-HA-bcd3UTR | This paper | N/A |
| pVAL22-Dmosk-Δ140-241-osk-HA-bcd3UTR | This paper | N/A |
| pVAL22-Dmosk-Δ401-606-HA-bcd3UTR | This paper | N/A |
| pVAL22-Gbosk2-90-Dmosk-HA-bcd3UTR | This paper | N/A |
| pVAL22-VcHlyU2-109-Dmosk-HA-bcd3UTR | This paper | N/A |
| pVAL22-DmLis2-86-Dmosk-HA-bcd3UTR | This paper | N/A |
| pVAL22-Dmosk-D155A-HA-bcd3UTR | This paper | N/A |
| pVAL22-Dmosk-S158R-HA-bcd3UTR | This paper | N/A |
| pVAL22-Dmosk-L165A-HA-bcd3UTR | This paper | N/A |
| pVAL22-Dmosk-T196S-HA-bcd3UTR | This paper | N/A |
| pVAL22-Dmosk-D197N-HA-bcd3UTR | This paper | N/A |
| pVAL22-Dmosk-S210P-HA-bcd3UTR | This paper | N/A |
| pVAL22-Dmosk-R215Q-HA-bcd3UTR | This paper | N/A |
| pVAL22-Dmosk-R215E-HA-bcd3UTR | This paper | N/A |
| pVAL22-Dmosk-T196E-D197V-R215I-HA-bcd3UTR | This paper | N/A |

| | | |
|---|---|---|
| *Vibrio cholera* transcription factor HlyU (residues 1 to 109) | DNASU Plasmid Repository | |
| pKNT25 | Euromedex | EUK001 |
| pKT25 | Euromedex | EUK001 |
| pUT18 | Euromedex | EUK001 |
| pUT18C | Euromedex | EUK001 |
| **Software and algorithms** | | |
| MAFFT | [5] | |
| PHENIX software package | [6–9] | |
| MODELLER | [10] | |
| HHpred webserver | [11] | |
| COOT | [12] | |
| PyMOL | [13] | |
| UNICORN software | GE Healthcare | |
| FIJI | [14] | |
| Python | Python Software Foundation | |
| FIJI macro code for microscopy image processing | This paper | https://github.com/Anastasiarep/LOTUS_mutant_analysis.git (commit ID 0de8b5a) |
| Python code for data analysis and figure generation | This paper | https://github.com/Anastasiarep/LOTUS_mutant_analysis.git (commit ID 0de8b5a) |
| Inkscape | Inkscape Project | |

#### SUPPLEMENTAL REFERENCES

1. Jeske, M., Bordi, M., Glatt, S., Müller, S., Rybin, V., Müller, C.W., and Ephrussi, A. (2015). The crystal structure of the Drosophila germline inducer Oskar identifies two domains with distinct Vasa helicase- and RNA-binding activities. Cell Rep. *12*, 587–598. https://doi.org/10.1016/j.celrep.2015.06.055.

2. Yang, N., Yu, Z., Hu, M., Wang, M., Lehmann, R., and Xu, R.-M. (2015). Structure of Drosophila Oskar reveals a novel RNA binding protein. Proc Natl Acad Sci U S A *112*, 11541–11546. https://doi.org/10.1073/pnas.1515568112.

3. Jeske, M., Müller, C.W., and Ephrussi, A. (2017). The LOTUS domain is a conserved DEAD-box RNA helicase regulator essential for the recruitment of Vasa to the germ plasm and nuage. Genes Dev. *31*, 939–952. https://doi.org/10.1101/gad.297051.117.

4. Ewen-Campen, B., Srouji, J.R., Schwager, E.E., and Extavour, C.G. (2012). oskar Predates the Evolution of Germ Plasm in Insects. Curr Biol *22*, 2278–2283. https://doi.org/10.1016/j.cub.2012.10.019.

5. Katoh, K., and Standley, D.M. (2013). MAFFT Multiple Sequence Alignment Software Version 7: Improvements in Performance and Usability. Mol. Biol. Evol. *30*, 772–780. https://doi.org/10.1093/molbev/mst010.

6. McCoy, A.J. (2006). Solving structures of protein complexes by molecular replacement with Phaser. Acta Crystallogr. Sect. D, Biol. Crystallogr. *63*, 32–41. https://doi.org/10.1107/s0907444906045975.

7. McCoy, A.J., Grosse-Kunstleve, R.W., Adams, P.D., Winn, M.D., Storoni, L.C., and Read, R.J. (2007). Phaser crystallographic software. J. Appl. Crystallogr. *40*, 658–674. https://doi.org/10.1107/s0021889807021206.

8. Adams, P.D., Grosse-Kunstleve, R.W., Hung, L., Ioerger, T.R., McCoy, A.J., Moriarty, N.W., Read, R.J., Sacchettini, J.C., Sauter, N.K., and Terwilliger, T.C. (2002). PHENIX: building new software for automated crystallographic structure determination. Acta Crystallogr. Sect. D *58*, 1948–1954. https://doi.org/10.1107/s0907444902016657.

9. Adams, P.D., Gopal, K., Grosse-Kunstleve, R.W., Hung, L., Ioerger, T.R., McCoy, A.J., Moriarty, N.W., Pai, R.K., Read, R.J., Romo, T.D., et al. (2004). Recent developments in the PHENIX software for automated crystallographic structure determination. J. Synchrotron Radiat. *11*, 53–55. https://doi.org/10.1107/s0909049503024130.

10. Webb, B., and Sali, A. (2016). Comparative Protein Structure Modeling Using MODELLER. Curr. Protoc. Bioinform. *54*, 5.6.1-5.6.37. https://doi.org/10.1002/cpbi.3.

11. Söding, J., Biegert, A., and Lupas, A.N. (2005). The HHpred interactive server for protein homology detection and structure prediction. Nucleic Acids Res *33*, W244–W248. https://doi.org/10.1093/nar/gki408.

12. Emsley, P., and Cowtan, K. (2004). Coot: model-building tools for molecular graphics. Acta Crystallogr. Sect. D: Biol. Crystallogr. *60*, 2126–2132. https://doi.org/10.1107/s0907444904019158.

13. Schrödinger, and LLC The PyMOL Molecular Graphics System, Version 3.0.

14. Schindelin, J., Arganda-Carreras, I., Frise, E., Kaynig, V., Longair, M., Pietzsch, T., Preibisch, S., Rueden, C., Saalfeld, S., Schmid, B., et al. (2012). Fiji: an open-source platform for biological-image analysis. Nat Methods *9*, 676–682. https://doi.org/10.1038/nmeth.2019.
